# Resistance Breaking in Root-knot Nematodes Carries a Fitness Cost Associated with Defective Feeding Site Development

**DOI:** 10.64898/2026.02.17.704924

**Authors:** Alison C. Blundell, Emma Shigekane-Kraft, Sławomir Janakowski, Mirosław Sobczak, Dadong Dai, Pallavi Shakya, Ching-Jung Lin, Valerie M. Williamson, Shahid Siddique

## Abstract

Root-knot nematodes (RKNs) cause an estimated 157 billion dollars in annual yield losses worldwide. In tomato, resistance to RKNs is conferred by the single dominant resistance gene *Mi-1*, which has been widely integrated into commercial cultivars. Prolonged and widespread use of *Mi-1* has led to the emergence of resistance-breaking populations in tomato fields worldwide; however, the consequences of resistance breaking for nematode performance on susceptible hosts remain poorly understood. Here, we compared infection outcomes of two closely related strains of *Meloidogyne javanica,* VW4 (Mi-1–avirulent; wild-type) and VW5 (Mi-1–virulent; resistance-breaking), on three susceptible hosts: tomato, cucumber, and rice. Across all hosts, VW5 produced significantly fewer eggs than VW4, revealing a fitness cost associated with Mi-1 virulence. Light and transmission electron microscopy of tomato and cucumber galls revealed impaired feeding site establishment by VW5. Consistent with these observations, transcriptomic profiling showed that VW5 infection induced weaker host transcriptional reprogramming and lacked gene expression signatures associated with effective suppression of plant defense responses. Together, these findings demonstrate that adaptation to Mi-1-mediated resistance incurs a fitness cost on susceptible plants and is accompanied by impaired feeding site formation and altered host reprogramming. Furthermore, these results establish VW4 and VW5 as a powerful resource for understanding nematode genes and pathways required for successful parasitism and feeding site development.

## Introduction

Plant-parasitic nematodes (PPNs) cause an estimated 157 billion U.S. dollars in annual crop losses worldwide (Kiontke and Fitch, 2013). Root-knot nematodes (RKNs; *Meloidogyne* spp.) are the most destructive group of PPNs and infect more than 3,000 plant species, including tomato, cucumber, soybean, and rice (Jones et al., 2013; Khan et. al., 2023; Blundell et al., 2026).

RKNs begin their life cycle as motile second-stage juveniles (J2s) that enter roots near the elongation zone and migrate toward the vascular cylinder. Upon reaching the vascular tissues, nematodes become sedentary and establish permanent feeding sites composed of modified host cells known as giant cells (GCs) (Wyss et al., 1992; Lin and Siddique). These GCs function as hypermetabolic sinks, and once established, nematodes remain sedentary and depend exclusively on them for nutrient acquisition throughout the remainder of their life cycle (Lin and Siddique, 2024). During feeding site initiation, maintenance, and development, RKNs extensively reprogram host tissues to support GC formation and function. Typically, five to seven GCs are induced per nematode and are characterized by cellular and nuclear hypertrophy, dense cytoplasm, an abundance of organelles, multinucleation, and an absence of a large central vacuole. These cells also develop extensive systems of cell wall ingrowths on walls facing newly differentiated xylem vessels, thus facilitating nutrient transport to the feeding site and nematode (Bartlem et. al., 2014). Without functional giant cells, nematodes are unable to feed and ultimately die (Hussey and Grundler, 1998).

The tomato resistance gene *Mi-1* (also referred to as *Mi-1.2*) has been incorporated into numerous tomato cultivars worldwide to manage RKNs. In some production systems, including processing tomatoes, it is estimated that more than 95% of commercial cultivars carry *Mi-1*. *Mi-1* confers resistance against several RKN species, including members of the *Meloidogyne incognita* group (*MIG; M. incognita, M. javanica,* and *M. arenaria*) and has also been reported to be effective against *M. luci* and *M. ethiopica* (Santos et al., 2020; Williamson, 1998; Roberts and Thomason, 1986). Notably, *Mi-1* does not prevent initial nematode penetration: J2s still enter roots and attempt to establish feeding sites. However, shortly after initial feeding cell selection, targeted host cells collapse, and a hypersensitive response (HR) is triggered at the attempted feeding site. HR can be detected as early as 12 hours post-infection, and nematodes either die from starvation or leave the root (Bleve-Zacheo et al., 1982; Paulson and Webster, 1972). Despite its efficacy, overreliance on *Mi-1* has resulted in the emergence of resistance-breaking RKN populations in tomato fields worldwide (Ploeg et al., 2023).

Previous studies have shown that resistance-breaking in plant parasitic nematodes can be associated with fitness costs on susceptible hosts. For example, reduced reproduction has been reported for resistance breaking strains of *M. incognita* on susceptible tomato and in some cases on pepper (Castagnone-Sereno et al., 2007; Djian-Caporalino et al., 2011; Ploeg et al., 2023). However, these studies generally lacked direct progenitors of the resistance breaking strains, making it difficult to determine whether reduced fitness reflected a true cost of resistance or simply differences in baseline virulence among genetically distinct populations.

Two closely related strains of *M. javanica,* VW4 (Mi-1–avirulent and unable to reproduce on Mi-1 tomato) and VW5 (derived from VW4 and Mi-1–virulent, capable of reproducing on resistant tomato) have been used as a model system to investigate mechanisms underlying *Mi-1* resistance-breaking (Gleason et al., 2008) (**Supp. Fig. 1**). Since VW5 was derived from a greenhouse culture of VW4 after only two generations of selection on resistant tomato, this strain pair also provides an opportunity to directly assess fitness costs associated with *Mi-1* virulence while minimizing confounding effects of unrelated genetic variation. As such, differences between VW4 and VW5 can be attributed more confidently to traits associated with Mi-1 virulence rather than inherent differences in overall pathogenicity.

In this study, we examined host responses and feeding site development using RNA-seq together with light and transmission electron microscopy of galled tissues. Our analyses show that VW5 is less fit on susceptible hosts and shows weaker host reprogramming and less developed feeding sites, consistent with a reproduction cost. Notably, this effect differed significantly with host. Together, these findings demonstrate that comparing VW4 and VW5 provides a useful approach to identify mechanisms required for successful nematode infection and feeding site development as well as clues to loss and gain of host range by these asexually-reproducing species.

## Methods

### Nematode Inoculation and Maintenance

*Meloidogyne javanica* populations were maintained on nematode-resistant *Solanum lycopersicum* resistant (cv. VFNT) and susceptible (cv. Momar Verte) plants (Wang et al., 2009). Seeds were germinated and transplanted under greenhouse conditions (26°C, nutrient irrigation via drainage 3 times a day for 3 minutes each) in 1-liter pots with sterile sand. To extract nematodes eggs, roots were rinsed to remove sand and cut into ∼1 cm segments. Root segments were shaken for 3 minutes in 10% [w/v] bleach (6% NaClO) and rinsed thoroughly with water over sieves (75 µm and 25 µm subsequently) until residual bleach was removed. Eggs were collected from the lower sieve into a 50 mL Falcon Tube. Following this, eggs were surface sterilized and set up for hatching as described in Liu and Williamson (2006). For infection assays, hatched J2s were counted and adjusted to 500 J2s/mL and each infected plant received 1 mL inoculum. All assays consisted of three independent experiments.

### Plant Material, Germination, and Growth

Experimental plant species included tomato (*S. lycopersicum* cv. Moneymaker and resistant cv. Motelle), cucumber (*Cucumis sativus* cv. Marketmore76), and rice (*Oryza sativa* cv. Kitaake). Seeds were surface sterilized prior to infection assays as described in Yimer et. al. (2023). Briefly, tomato and rice seeds were germinated on moist filter paper in darkness at 28°C for 5-7 days, whereas cucumber seeds were directly planted into pots. Plants used for greenhouse assays were grown in 1-liter Styrofoam pots filled with sterile sand under the same conditions as described above for nematode culture maintenance.

### Nematode Infection Assays

Seedlings were inoculated with either VW4 or VW5 J2s. For each assay, three independent experimental replicates were conducted. For nematode eggs collection, roots were harvested at 35 days post inoculation (dpi) and aboveground tissues were discarded. Fresh root weight was recorded in grams. For infection assays, roots were not cut prior to egg collection to allow subsequent staining when required. Collected eggs were counted for all plant cultivars tested. For tomato, stained roots were also used to count the numbers of females and galls.

### Root Staining and Nematode Visualization

Following egg collection, intact tomato roots were stained in Acid Fuchsin (ThermoFisher Scientific, Waltham, MA, USA: 227905000) Roots were immersed in boiling stain for 3 minutes with constant agitation, rinsed thoroughly with water, and stored in glycerol until observation (Byrd et al., 1983).

### Quantification of Nematode Life Stages and Galls

Nematodes and galls were counted using a Leica S8 APO dissecting microscope (Leica Microsystems, Wetzlar, Germany). Each nematode was counted as a single individual. Galls were dissected as needed to ensure accurate nematode counts. Each data point represents counts per individual plant.

### Invasion Assays

Tomato (cv. Moneymaker) seedlings were germinated as described above. Seedlings were transplanted in a controlled growth room in sterilized sand with polymer mixture as described in Yimer et al. (2023). Plants were grown in 4-inch plastic pots and watered using half strength Hoagland’s solution 3 times a week (Reversat et al., 1999). Plants were maintained under photoperiod 12-hr light: 12-hr dark cycle. Plants were harvested at 3 dpi and total J2s were counted per root system.

### Infection Assays for Transcriptional Analysis

Due to visible abnormalities in gall morphology, cucumber (cv. Marketmore76) plants were used for mRNA-Seq analysis and microscopy. Plants were grown under controlled growth room conditions as described above. Galls and uninoculated roots were collected at 7 and 28 dpi following infection by VW4 or VW5. Upon dissection, excised galls were flash frozen in liquid nitrogen. Samples were randomly pooled into three sample batches and stored in the -80°C until RNA extraction.

### RNA Extraction and Sequencing

RNA was extracted using the RNeasy Mini Kit (QIAGEN, Hilden, Germany: 74106). Samples were ground using metal beads and a tissue homogenizer. RNA concentrations were measured using a NanoDrop spectrophotometer (ThermoFisher Scientific: 13-400-526) and adjusted to needed values for sequencing. RNA samples were submitted to Novogene (Novogene Corporation Inc., Sacramento, CA, USA). Samples were checked for quality and ran on a NovaSeq X Plus 25B instrument and complied into a mRNA library preparation (poly A enrichment) using the ABClonal mRNA-Seq Library Prep Kit for Illumina (Catalog: RK20302).

### RNAseq Processing Differential Expression Analysis

The raw data was filtered and trimmed to obtain clean reads (Dong et al., 2026). The cleaned reads were then mapped to the cucumber reference genome Chinese long version 4 (Clv4) (Guan et al., 2024) using HISAT2 (Kim et al., 2019). Similarly, nematode reads were mapped to *M. javanica* (VW4) reference genome (Winter et al., 2024). After mapping, the raw counts for each gene. Differential expression gees (DEGs) analyses were performed using DESeq2 (Love et al., 2014). Raw read counts were modeled using a negative binomial distribution in DESeq2, and differential expression was assessed using the Wald test. P-values were adjusted for multiple testing using the Benjamini-Hochber procedure. Genes with an adjusted p-value < 0.05 and l log2fold change l >1 were considered DEGs. To generate volcano plots, the adjusted p-values were ranked from smallest to largest. For data analysis, ggplot and dplyr packages were used where the x-axis = log_2_(Fold change) and the y-axis = -log10(adjusted p-value). (GitHub https://github.com/Siddique-Lab/Resistance_Breaking_Manuscrip<u>^t^</u>). Venn diagrams were generated using GraphPad Prism v.10.0.0.

### GO Analysis and Transcriptional Pathways Analysis

For functional analysis eggNOG-mapper (Huerta-Cepas et al., 2017) was used for KEGG and Gene ontology (GO) annotation, pfam_scan.pl (Mistry et al., 2021) was used to predict the Pfam conserved domain. GO-enrichment analysis was performed using TBtools-II (chen et al., 2020) in R (v.4.5.2) and Rstudio software (v.2025.09.2+401) followed by using R studio package ggplot to identify the top 15 most significant gene functions for each data set. To identify the top 15 most significant gene functions for each data set, gene ratios were calculated as a ratio of genes counts in the selected set (adjusted p-value) and gene counts in the background (number of significant genes with the same function). For all data processed, adjusted p-values were labeled at a significant threshold of *p < 0.05* and l log_2_(fold change) l < 1.

### Sampling of Galled Tissues for Microscopy

Susceptible (cv. Moneymaker) and resistance (cv. Motelle) tomato along with cucumber (cv. Marketmore76) plants were grown for 21 days before inoculation with J2s of *M. javanica* as described above. Collected samples consisted of galled root segments and uninoculated root segments at comparable locations and were collected at 7 and 28 dpi under a dissection microscope as mentioned in RNA-Seq assays. Root segments were immediately transferred into a modified Karnovsky’s fixative solution composed of 2% [w/v] paraformaldehyde and 2% [w/v] glutaraldehyde dissolved in 50 mM sodium cacodylate buffer, pH = 7.2. The samples were incubated for 2 hours and thereafter the fixative was replaced with 50 mM cacodylic buffer (pH = 7.2) and incubated for 10 mins. The washing steps were repeated 4 times and the samples were stored in the last buffer bath until further processing. Around 10 galls were placed per 1.5 mL tube. Gall and root segments were between 6-7 mm in size.

### Gall Sectioning and Imaging

The samples were post-fixed in a 2% [w/v] aqueous solution of osmium tetroxide (OsO_4_) at room temperature in darkness for 2 hours and rinsed with cacodylic buffer as described above. Following this, samples were dehydrated in a graded series of ethanol solutions (10% [v/v] increments) for 15 minutes each. The final dehydration was performed with pure ethanol (with 3 baths over 60 minutes in total). Ethanol was then removed and substituted with pure propylene oxide (3 baths for 20 minutes each) before sample infiltration with mixtures (4:1, 2:1, 1:1, 1:2, 1:3; 1:4 [v/v]) of propylene oxide and EPON-type epoxy resin (Agar Scientific, Stansted, UK) for 2 hours each. Samples were then incubated in pure resin for 24 hours (replacing resin every 6 hours). After this, the samples were transferred into flat embedding molds filled with pure resin and positioned for transverse sectioning. Finally, the resin was cured at 65°C for 12 hours.

Serial sections for light microscopy examinations (3 µm thickness) were produced on glass knives using a Leica RM2165 microtome (Leica Microsystems). Sections were stained in 0.1% [w/v] aqueous solution of Toluidine Blue in 50 mM phosphate buffer (pH = 6.9) at 65°C for 3 minutes. They were examined under an Olympus AX70 ‘Provis’ light microscope (Olympus, Tokyo, Japan) in the bright-field mode. Digital images were taken with an Olympus UC90 digital camera (Olympus). They were resized, cropped and equalized for similar contrast and brightness using Adobe Photoshop image processing software (Adobe Inc., San Jose, CA, USA).

Sections for transmission electron microscopy were taken at defined places based on analyses of light microscopy images. Ultra-thin (80 nm thickness) sections were cut on a Leica Ultracut E ultramicrotome (Leica) equipped with a diamond knife (DiATOME, Nidau, Switzerland). The sections were collected on formvar coated, single slot copper grids (Agar). Prior to examinations sections were counter-stained in saturated solution of uranyl acetate in 70% [v/v] methanol and 2.5% [w/v] aqueous solution of lead citrate. After air-drying the sections were examined under a FEI268d ‘Morgagni’ transmission electron microscope (FEI Comp., Hillsboro, OR, USA) operating at 80 kV and equipped with a SIS-Olympus ‘Morada’ digital camera (Olympus) with 16 MPix resolution. Digital images were resized and equalized for similar contrast and brightness using Adobe Photoshop.

### Statistical Analysis

For all infection assays, GraphPad Prism (https://www.graphpad.com/) was utilized for statistical analysis. Outliers were removed using GraphPad Prism ROUT method with Q=10%, cleaned data was used and two tailed unpaired student’s t-test with a 95% confidence interval, assuming Gaussian distribution.

### Data Availability

The transcriptomic datasets generated in this study have been deposited in the NCBI database under BioSample accession number SAMN55364028 (https://www.ncbi.nlm.nih.gov/biosample/55364028 All code and command-line workflows used for RNA-seq analyses are publicly available at GitHub: https://github.com/Siddique-Lab/Resistance_Breaking_Manuscript.

## Results

### VW5 Shows Reduced Reproduction Compared to VW4 on Susceptible Tomato

To evaluate whether *Mi-1*-virulence is associated with a fitness cost on susceptible hosts, we compared parasitism parameters of *M. javanica* VW4 and VW5 on susceptible tomato (cv. Moneymaker). Plants were harvested at 35 dpi and the average numbers of eggs, females, galls, nematodes per root (all life stages), and fresh root weight were determined. Although several parameters showed differences among independent trials, only lower egg production in VW5 compared to VW4 was consistent across all assays (**Figure 1A**–**1D** and **Supp. Table 1**). No significant difference in root weight was observed between genotypes (**Supp. Fig. 2A**). Overall, these data identified a reduction in egg production as the most consistent and robust phenotype distinguishing VW5 from VW4 on susceptible tomato.

**Figure 1:**
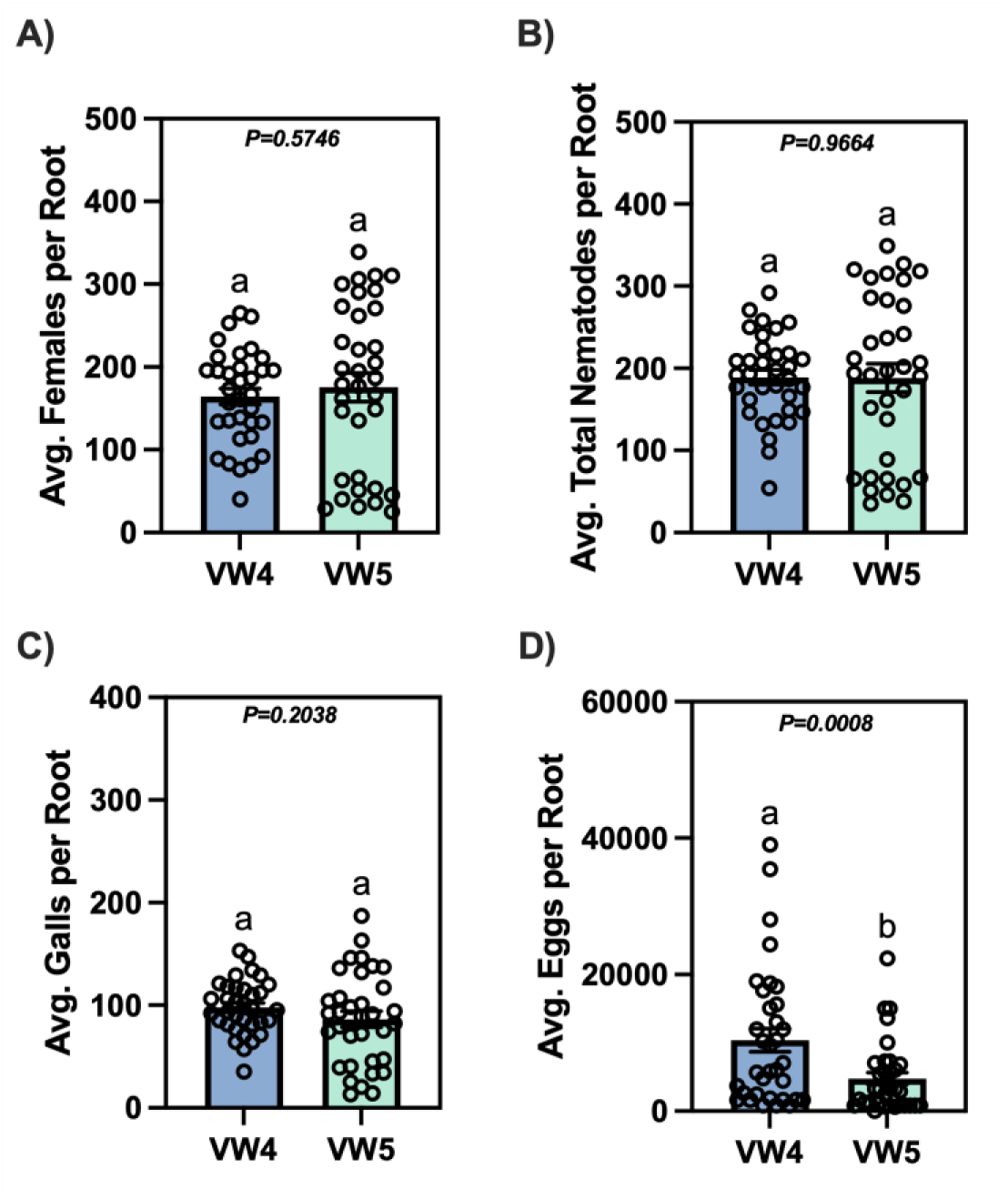
VW5 shows reduced reproduction compared with VW4 on susceptible tomato. Infection assays were performed on susceptible tomato cv. moneymaker using *M. javanica* VW4 (wildtype, Mi-1–avirulent) and VW5 (Mi-1–virulent, resistance-breaking). Roots were harvested at 35 dpi and following metrics were evaluated: (A) Average number of females per root system, (B) Average number of total nematodes (all developmental stages) per root system, (C) Average number of galls per root system, and (D) Average number of eggs per root system. Each data point represents an individual root system, and bars indicate means ± standard error of the mean (SEM). Data is pooled from three independent biological experiments (n = 35 roots per treatment). Statistical significance was determined using unpaired two-tailed Student’s t-tests (p < 0.05). Different letters indicate statistically significant differences between VW4 and VW5.

### VW4 and VW5 Invade Tomato Roots at Comparable Rates

Considering that VW5 exhibited reduced egg production, we next asked whether this phenotype could be due to impaired nematode penetration of roots. Since total nematode counts within roots did not significantly differ between VW4 and VW5 at 35 dpi (**Figure 1B**), we examined early root penetration directly to see if penetration was delayed. J2s were inoculated onto susceptible tomato (cv. Moneymaker) roots and were harvested at 3 dpi. Roots were cleaned and stained with Acid Fuchsin and the number of J2s were counted per root system (**Figures 2A** and **2B**). No significant difference was found in the number of invading J2s between VW4 and VW5 at 3 dpi, indicating that both strains enter susceptible tomato roots at comparable levels. Together, this data indicates that reduced eggs production in VW5 is not due to delayed or reduced penetration but likely reflects defects in the ability of VW5 to develop into an egg laying female.

**Figure 2:**
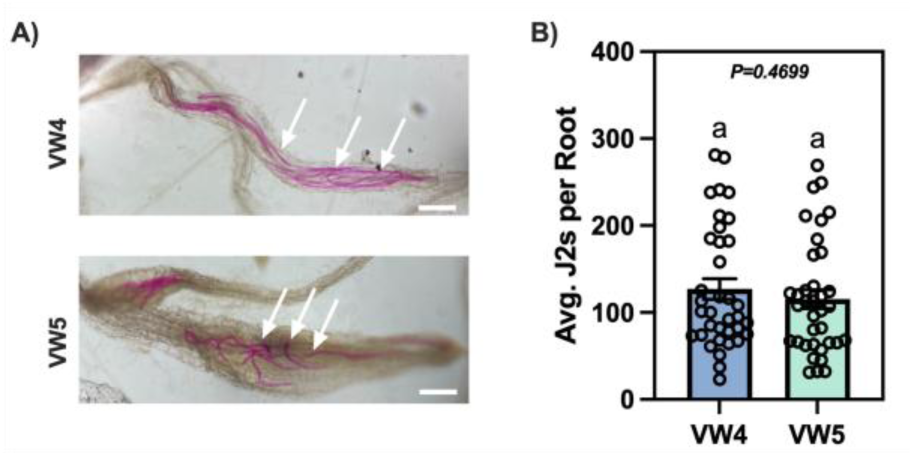
VW4 and VW5 invade susceptible tomato roots at comparable rates. (A) Representative images of Acid Fuchsin–stained tomato roots showing VW4 and VW5 J2s within root tissues. (B) Quantification of average number of J2s per root for VW4 and VW5. Each data point represents an individual root system. Arrows indicate J2s in roots. Bars indicate mean ± standard error of the mean (SEM). Statistical significance was assessed using an unpaired Student’s t-test; no significant difference was detected (n = 36 roots per treatment). Scale bars = 350 µm.

### VW5 Exhibits Reduced Egg Production Compared to VW4 on Cucumber and Rice

To determine whether the reduced reproduction of VW5 occurred in other hosts besides tomato, we tested the VW4 and VW5 strains on additional susceptible hosts: cucumber (cv. Marketmore76) and rice (cv. Kitaake). Cucurbits are common rotation crops for tomato and rice was included as a monocot host. Plants were harvested at 35 dpi, and fresh root weight along with nematode egg counts were quantified. As observed on tomato, VW5 produced significantly fewer eggs than VW4 on both cucumber and rice. Notably, the reduction in egg production was more pronounced with a 90% reduction in cucumber and an 88% reduction in rice than that found for susceptible tomato (58% reduction) (**Figures 3A** and **3B**). Fresh root weight was recorded for cucumber and rice, and we found that there was a significant difference for cucumber root weight between VW4 and VW5 infection. However, this may be explained by a significantly smaller population size for cucumber compared to tomato and rice (**Supp. Figures 2B** and **2C**).

**Figure 3:**
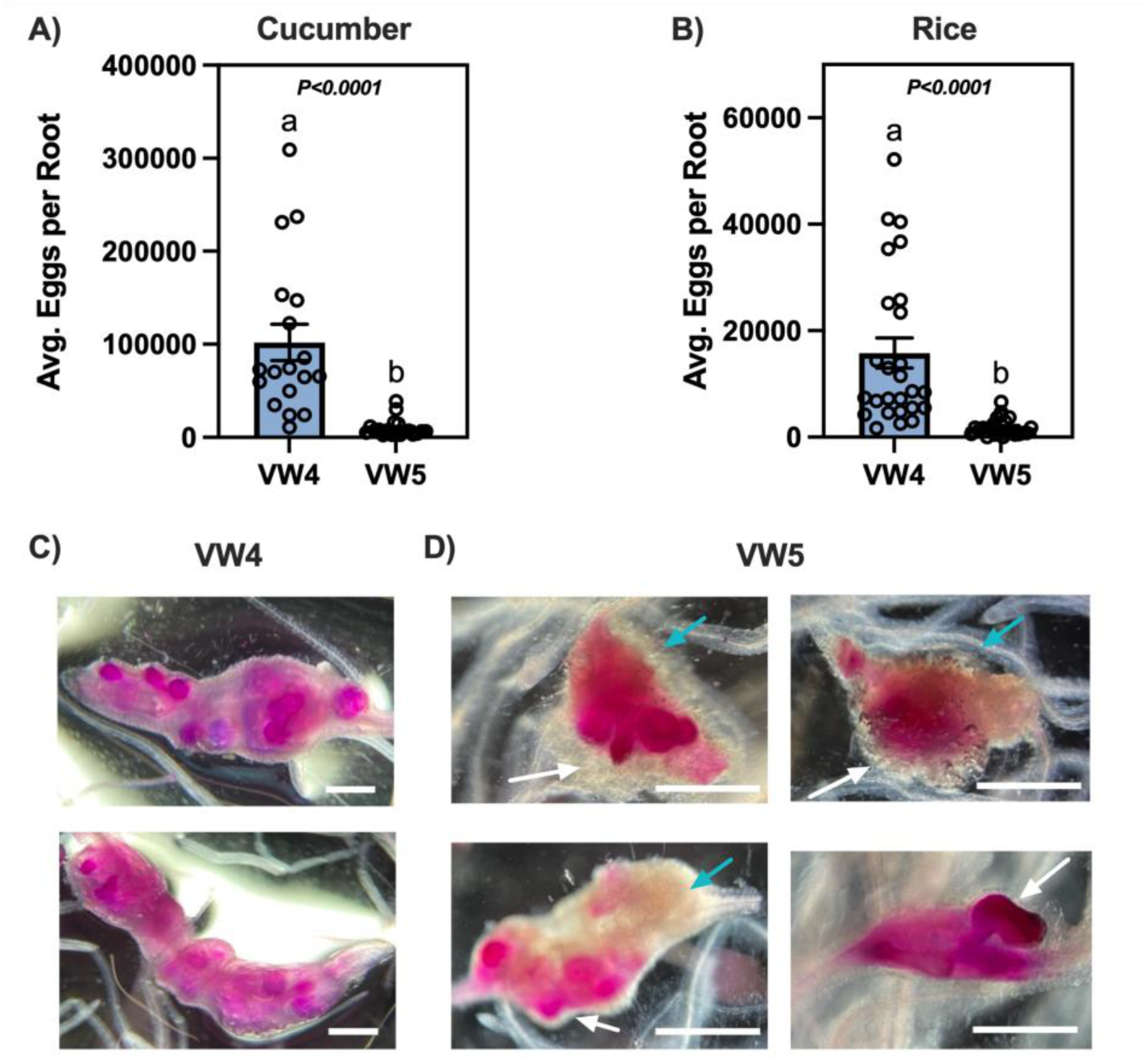
VW5 shows reduced egg production on cucumber and rice and induces abnormal gall morphology on cucumber. (A) Average egg numbers per cucumber root; (B) Average egg numbers per rice root. Each data point represents an individual root system. Bars indicate means ± standard error of the mean (SEM). Statistical significance was determined using unpaired Student’s t-tests, with p values shown; different letters indicate statistically significant differences (p < 0.05). Data are pooled from three independent biological experiments (cucumber, n = 18; rice, n = 26). (C and D) Cucumber roots were stained with Acid Fuchsin to visualize nematodes and gall morphology. Representative images show galls induced by VW4 (C) and VW5 (D). VW5 female nematodes are indicated with white arrows and dense cellular tissue is indicated in blue arrow. Scale bars = 350 µm.

To assess gall morphology, we stained a subset of cucumber roots with Acid Fuchsin for nematode visualization. VW5-infected cucumber roots displayed pronounced abnormalities in gall morphology compared with VW4 (**Figure 3C** and **3D**). At 35 dpi, galls produced by VW4 contained globular females embedded within the galled tissue with nematode heads aligned along vascular tissue (**Figure 3C**). Intriguingly, VW5 females were frequently found at the periphery of the gall, singular and often distorted females, and galls contained dense callus-like tissue in regions where no females were present (**Figure 3D**). Rice roots were also stained; however, difficulty visualizing the nematodes within the roots precluded further morphological analysis.

### VW5 Shows Impaired Feeding Site Development

To further investigate differences in gall morphology between VW4 and VW5, we examined transverse sections of galls and uninoculated roots by light and transmission electron microscopy. Galls were collected at 7 and 28 dpi from resistant tomato (cv. Motelle), susceptible tomato (cv. Moneymaker), and cucumber (cv. Marketmore76) infected with VW4 or VW5.

Uninoculated tomato roots displayed typical root anatomy, including a single rhizodermis (epidermis) cell layer, a cortex composed of several cell layers, an endodermis, and a diarchal vascular cylinder with two primary xylem and two primary phloem bundles surrounded by a single layer of pericyclic cells (**Supp. Figures 3A** and **3B**). Similarly, uninoculated cucumber roots showed parallel organization with only slight differences in endodermal cell morphology and a triarchic vascular cylinder with a central metaxylem vessel in the center (**Supp. Figure 3C**). Above the root-hairs, the formation of secondary thickening commenced (**Supp. Figures 3D-3F**). The rhizodermis deteriorated and was replaced by the exodermis, and secondary phloem and xylem elements started to differentiate between primary xylem and phloem bundles. No formation of secondary cover tissue (periderm) was observed.

Cross sections of galled cucumber roots were taken at the nematode head region and were only selected for comparisons of feeding site anatomy. VW4 and VW5 feeding sites in cucumber displayed slower GC selection and development than in tomato, especially in the case of VW5 (**Figures 4A**–**4D** vs **Supp. Figures 4A-4C**). At For both nematode strains at 7 dpi, many invaded J2s had not established feeding sites (**Figures 4A** and **4B**). Presumably, the invasion took place in the root-tip regions and induced extensive divisions within the vascular cylinder cells without differentiation into xylem vessels. However, some successful VW4 juveniles established groups of 2-3 clearly recognizable GCs (**Figure 4C**) whereas most VW5 juveniles were found surrounded by a ring of only enlarged cells with central vacuoles (**Figure 4D**).

**Figure 4:**
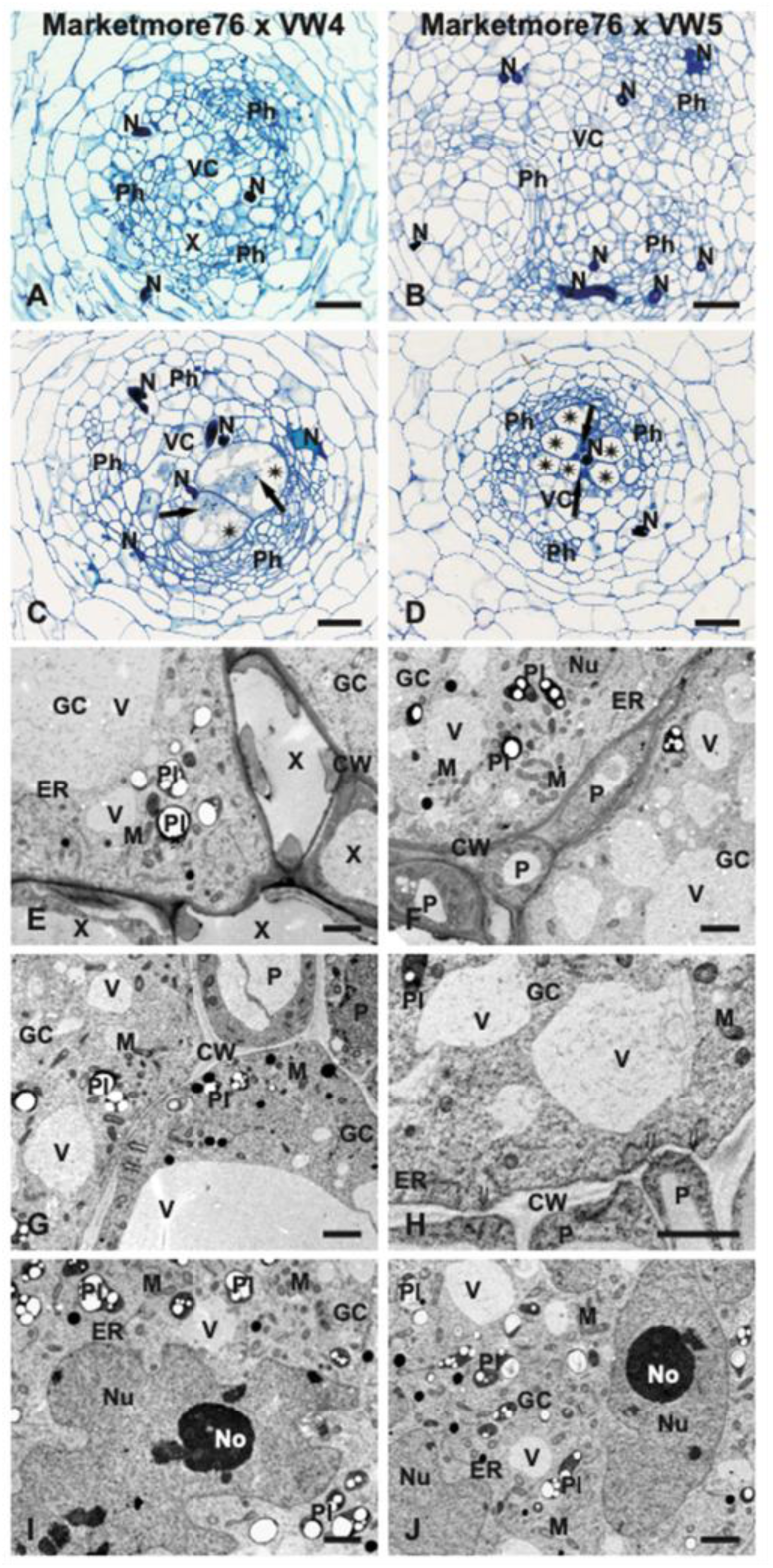
Anatomy and ultrastructure of giant cells developed in cucumber roots at 7 dpi. Light (A-D) and transmission electron microscopy (E-J) images of transverse sections of roots infected with VW4 strain (A, C, E, G and I) or VW5 strain (B, D, F, H and J). Asterisks indicate giant cells (C and D). Arrows point to giant cell nuclei (C and D). Double tails arrows indicate plasmodesmata in thin fragments of giant cell walls (G and H). Abbreviations: CW, cell wall; ER, endoplasmic reticulum; GC, giant cell; M, mitochondrion; N, nematode; No, nucleolus; Nu, nucleus; P, parenchymatic cell; Ph, phloem; Pl, plastid; X, xylem vessel; V, vacuole. Scale bars: 50 µm (A-D) and 2 µm (E-J).

In the case of VW5, the GCs or “enlarged cells” were usually surrounded by parenchymatic cells (**Figures 4F** and **4H**) and only few xylem vessels contained protoplasts (**Figures 5F**). In contrast, VW4-induced GCs were frequently surrounded by fully differentiated xylem vessels (**Figure 4E** and **Figure 5E**) and no cell wall ingrowths were observed at 7 dpi but were present at 28 dpi GCs (**Figures 4E**–**4H**, and **Figures 5E**, **5F**, **5I**, and **5J**). Some parts of the cell walls between the GCs and the neighboring parenchymatic cells remained thin and pierced with plasmodesmata at both time points (**Figures 4G** and **4H**). Protoplasts of GCs contained vacuoles of different sizes at 7 dpi (**Figures 4C**–**4J**), but only small ones were present in GCs with non-degraded protoplasts at 28 dpi (**Figures 5E**–**5H**). The cytoplasm contained numerous mitochondria, plastids, and relatively few elements of the endoplasmic reticulum (**Figures 4E**–**4J** and **Figures 5E**–**5I**). Plastids and mitochondria had typical round or rod-shaped outlines, and no morphologically modified organelles were observed in cucumber but often occurred in GCs induced in tomato roots. Starch grains were abundantly present in plastids at 7 dpi (**Figures 4E**–**4J**), but at 28 dpi, they only appeared in high numbers for GCs induced by VW5 (**Figures 5E**–**5J**).

**Figure 5:**
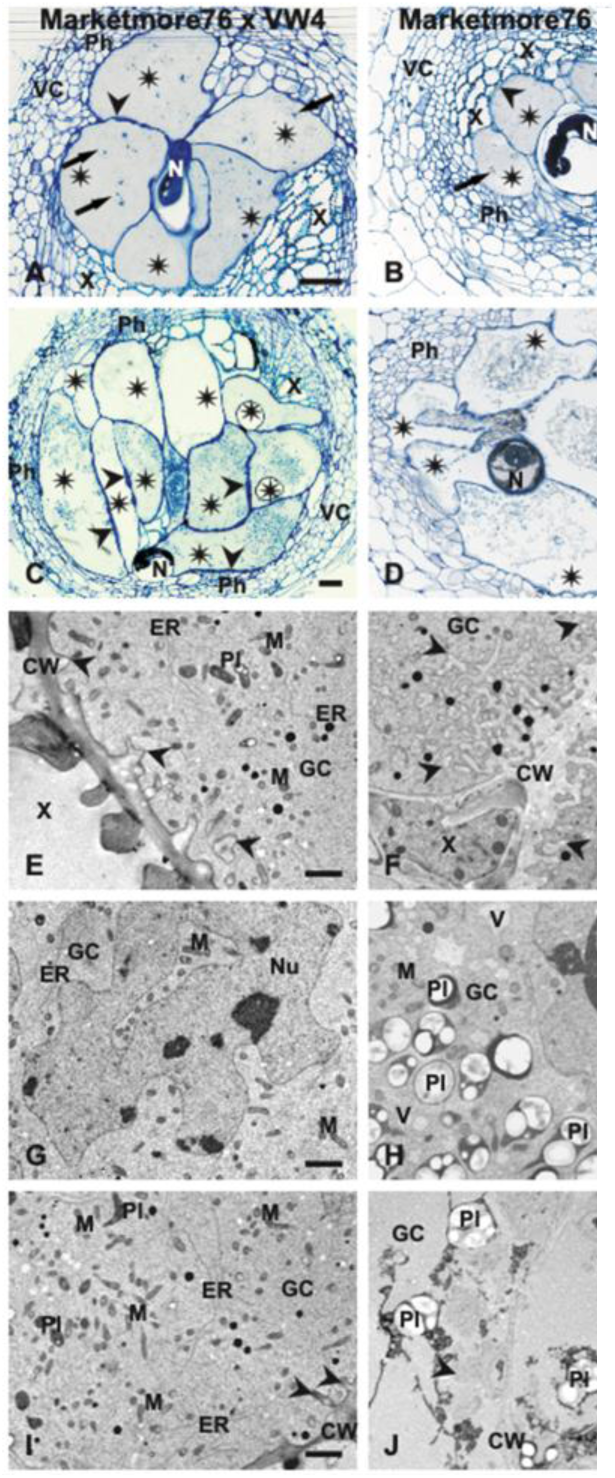
Anatomy and ultrastructure of giant cells developed in cucumber roots at 28 dpi. Light (A-D) and transmission electron microscopy (E-J) images of transverse sections of roots infected with VW4 strain (A, C, E, G and I) or VW5 strain (B, D, F, H, and J). Asterisks indicate giant cells (A-D). Arrows point to giant cell nuclei (A and B). Arrowheads indicate systems of cell wall ingrowths (A-F, I and J). Abbreviations: CW, cell wall; ER, endoplasmic reticulum; GC, giant cell; M, mitochondrion; N, nematode; No, nucleolus; Nu, nucleus; Ph, phloem; Pl, plastid; X, xylem vessel; V, vacuole. Scale bars: 50 µm (A-D) and 2 µm (E-J).

In general, the GC nuclei were strongly hypertrophied and contained electron-dense nucleoplasm and enlarged nucleoli for both timepoints and nematode strains (**Figures 4I** and **4J**, and **Figures 5G** and **5H**). Additionally, GCs became strongly amoeboid in outlines except for nuclei of GCs induced by VW5 at 7 dpi, which were hypertrophied but remained relatively round (**Figures 4F** and **4J**).

At 28 dpi, two different patterns of feeding site anatomy could be discriminated between the two strains (**Figures 5A**–**5D**). In VW5 feeding sites, some were composed of 4-6 strongly hypertrophied GCs surrounding the nematode head and with extensive interfaces to the secondary xylem vessels (**Figures 5A** and **5B**). The cell walls of the GCs were slightly thickened, and systems of cell wall ingrowths were weakly developed (**Figures 5A**, **5B**, **5E**, and **5F**). If deposited, the cell wall ingrowths were usually formed at walls facing xylem vessels (**Figures 5E**, **5F**, and **5I**). GCs cytoplasm contained only small vacuoles, numerous mitochondria and plastids of typical round or elongated outlines (**Figures 5E**–**5I**). However, in some cases, feeding sites induced by both nematode strains contained additional GCs that did not make contact with the nematode head (**Figures 5C** and **5D**). Such GCs contained usually poorly stainable and granular protoplast being degraded in the case of VW4 (**Figure 5C** and **5I**) or degraded in the case of VW5 (**Figure 5D** and **5J**).

VW4 and VW5 feeding sites in tomato displayed similar results, however feeding site initiation and development was more progressed than observed in cucumber galls. Cross sections revealed that GCs were consistently located within the vascular cylinder adjacent to conductive elements of xylem and phloem (**Supp. Figures 4A-4C** and **5A-5C**). Infected roots showed marked proliferation of vascular cylinder cells relative to uninfected roots (**Supp. Figures 4A-4C** vs **Supp. Figures 3A** and **3B**). During VW5 parasitism of susceptible tomato, GCs were frequently displaced toward the periphery of expanded vascular cylinders, with limited contact surface to the conductive elements (**Supp. Figure 4E**).

Clear differences in feeding sites were seen at 7 dpi. Typical feeding sites induced in Moneymaker roots by VW4 juveniles contained 4-6 strongly hypertrophied GCs surrounding the nematode head at 7 dpi (**Supp. Figure 4A**) and the number of GCs rose to 6-8 at 28 dpi (**Supp. Figure 5A**). In contrast, VW5 infections on both Moneymaker and Motelle frequently contained more than 6 GCs, including additional GCs not in direct contact with the nematode body, giving the impression of a second GCs ring (**Supp. Figures 4B, 4C** and **4E**). Despite higher GC numbers, VW5-induced GCs were usually less hypertrophied and rather thin-walled than those induced by VW4 at both 7 and 28 dpi (**Supp. Figures 4A-4C and 4E** and **Supp. Figures 5A-5C**).

VW4-induced GCs exhibited extensive cell wall thickening and branched wall ingrowths between GCs and adjacent xylem vessels and sieve tubes (**Figure 6** and **Supp. Figures 4A, 4D, 4G** and **4J**), which became more pronounced at 28 dpi (**Figure 7** and **Supp. Figures 5A, 5D** and **5G**). In contrast, ingrowths were reduced or absent in VW5-induced GCs, particularly in susceptible Moneymaker roots (**Figure 6** and **Supp. Figures 4B, 4C, 4F, 4H, and 4L**; **Figure 7** and **Supp. Figures 5B, 5C, 5E** and **5F**). Thin parts of cell walls containing plasmodesmata were frequently observed between GCs in VW4 and VW5 infections at both time points (**Supp. Figures 4G** and **4L**, and **Supp. Figures 5G** and **5I**).

**Figure 6:**
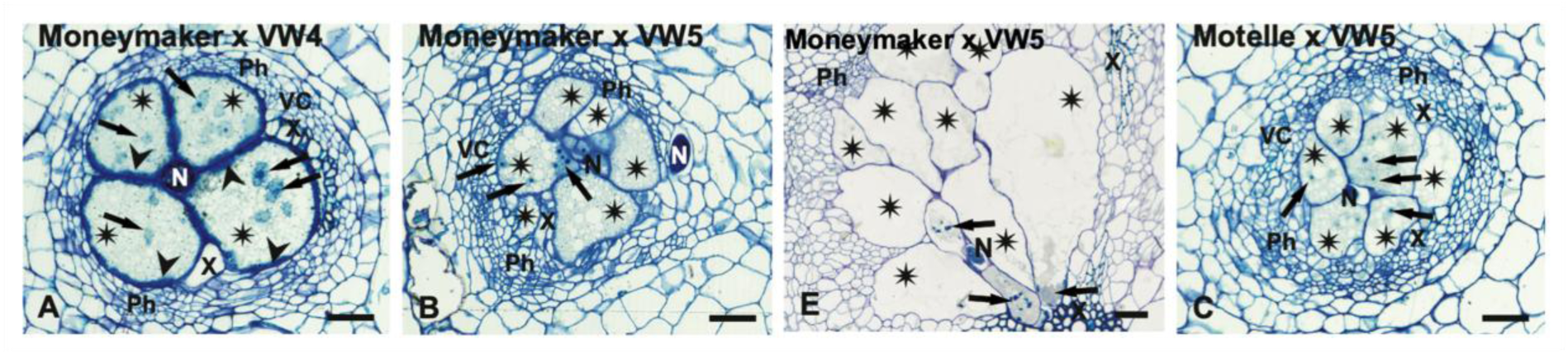
Anatomy and ultrastructure of giant cells developed in tomato roots at 7 dpi. Light (A-C and E) images of transverse sections of susceptible cv. Moneymaker (A, B, and E) and resistant cv. Motelle (C) infected with avirulent VW4 strain (A) or virulent VW5 strain (B, C, and E). Asterisks indicate giant cells (A-C and E). Arrows point to giant cell nuclei (A-C and E). Arrowheads indicate systems of cell wall ingrowths (A and D). Scale bars: 50 µm (A-C and E). Abbreviations: N, nematode; Ph, phloem; X, xylem vessel; and VC, vacuole. Additional microscopy images for tomato at 7 dpi including TEM images can be found in Supp. Figure 4.

**Figure 7:**
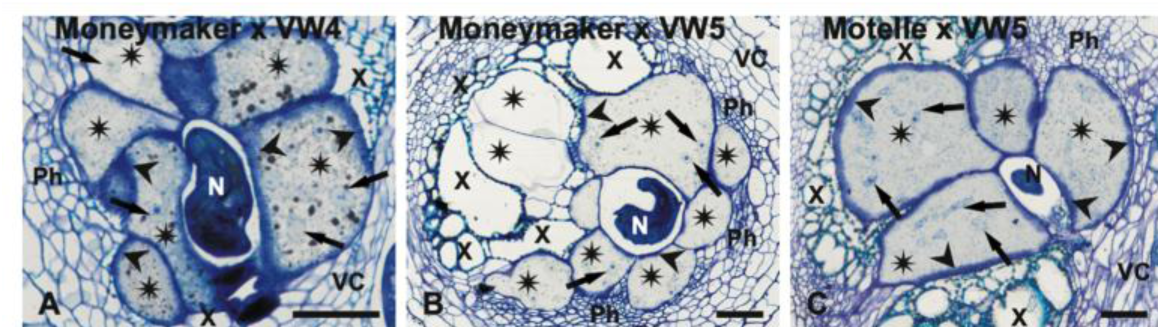
Anatomy and ultrastructure of giant cells developed in tomato roots at 28 dpi. Light (A-C) of susceptible cv. Moneymaker (A and B) and resistant cv. Motelle (C) infected with avirulent VW4 strain (A) or virulent VW5 strain (B and C). Scale bars: 50 µm (A-C). Abbreviations: N, nematode; Ph, phloem; X, xylem vessel; and VC, vacuole. Additional microscopy images for tomato at 28 dpi including TEM images can be found in Supp. Figure 5.

At the ultrastructural level, protoplasts of VW4-induced GCs stained strongly with Toluidine Blue and their cytoplasm was uniformly electron dense at both 7 and 28 dpi. The cytoplasm was rich in organelles, including mitochondria, plastids, limited Endoplasmic reticulum (ER) structures, hypertrophied nuclei, and numerous in small vacuoles which were evenly distributed inside the GCs (**Supp. Figures 4D, 4J** and **4M** and **Supp. Figures 5J** and **5M**). Most of the mitochondria were not morphologically changed, however, at 28 dpi many of them acquired elongated, circular or cup-like shapes (**Supp. Figures 4D** and **4J** and **Supp. Figures 5G, 5J** and **5M**). Plastids appeared numerously and rarely contained small starch grains (**Supp. Figures 4D, 4G, 4J** and **4M,** and **Supp. Figures 5G, 5J** and **5M**). ER cisternae were rare and usually formed short structures at both timepoints (**Supp. Figures 4D, 4G** and **4J,** and **Supp. Figures 5D** and **5G**). GC nuclei were strongly hypertrophied, amoeboid in outlines, and contained uniformly electron dense nucleoplasm with prominent nucleoli at 7 and 28 dpi (**Supp. Figures 4J** and **4M** and **Supp. Figure 5M**).

In contrast, the cytoplasm of GCs induced by VW5 in Moneymaker was frequently electron-translucent and usually contained large vacuoles resembling the central vacuole (**Supp. Figures 4E, 4K** and **4N**). Nevertheless, at 28 dpi these vacuoles were reduced in size and two zones of cytoplasm could be discriminated: the opaquer layer next to the cell walls which contained the bulk of organelles and the transparent central part containing only vacuoles of different shapes and sizes (**Supp. Figures 5B, 5H** and **5K**). The mitochondria and plastids were less numerous (**Supp. Figures 4H, 4K** and **4N,** and **Supp. Figures 5E, 5H, 5K** and **5N**) and were not changed morphologically at any time point. Additionally, plastids usually contained small starch grains (**Supp. Figures 4H** and **Supp. Figure 5E**). ER cisternae were more numerous than observed in VW4 and were often arranged in circular structures in 7 dpi and 28 dpi GCs (**Supp. Figures 4H** and **4K** and **Supp. Figures 5E** and **5K**). In contrast, nuclei were slightly hypertrophied and appeared as round forms with small bulges at both time points, when compared to amoeboid observations in GCs induced by VW4 (**Supp. Figures 4J, 4K** an**d 4M** and **Supp. Figures 5H, 5M** and **5N**).

VW5-induced GCs in resistant Motelle showed resemblance as observed in VW4-induced GCs, especially at 28 dpi (**Figure 6** and **Supp. Figures 4A, 4C, 4D** and **4F** and **Figure 7** and **Supp. Figures 5C, 5D** and **5F**). The GCs cytoplasm displayed similar opacity and contained numerous small vacuoles (**Supp. Figures 4F, 4I** and **4L,** and **Supp. Figures 5F, 5I, 5L,** and **5O**). In addition, they contained abundant mitochondria, which often acquired elongated or ring-like shapes at 28 dpi (**Supp. Figures 5F, 5I,** and **5L**). Many plastids contained large starch granules (**Supp. Figures 4F, 4I** and **4L, and Supp. Figures 5L** and **5O**). Similarly, nuclei were strongly hypertrophied, amoeboid in shape, and contained prominent nucleoli at 7 and 28 dpi (**Supp. Figure 4O** and **Supp. Figure 5O**). Together, these light and transmission electron microscopy analyses demonstrate that VW5 establishes structurally compromised feeding sites during infection on susceptible hosts, characterized by reduced GCs hypertrophy, limited vascular connectivity, and altered organization of GCs protoplast, which likely leads to premature degradation of its feeding sites, restricts its nutrient supply and contributes to their reduced reproduction.

### Transcriptomes of Cucumber Galls Reveal Weaker Host Reprogramming by VW5 Compared to VW4 at Early Infection

To investigate host transcriptional responses associated with the observed feeding site defects and reduced reproduction, we performed RNA-seq analysis with cucumber galls at 7 and 28 dpi with VW4 or VW5. Principal component analysis (PCA) showed clear separation between infected and uninoculated samples along PC1 at both time points, indicating strong transcriptional reprogramming by the nematodes following infection (**Figures 8A** and **8B**). In addition, VW4 and VW5 samples clustered closer to each other than to uninoculated controls, suggesting a broadly similar host response yet with discernible strain-specific differences.

**Figure 8:**
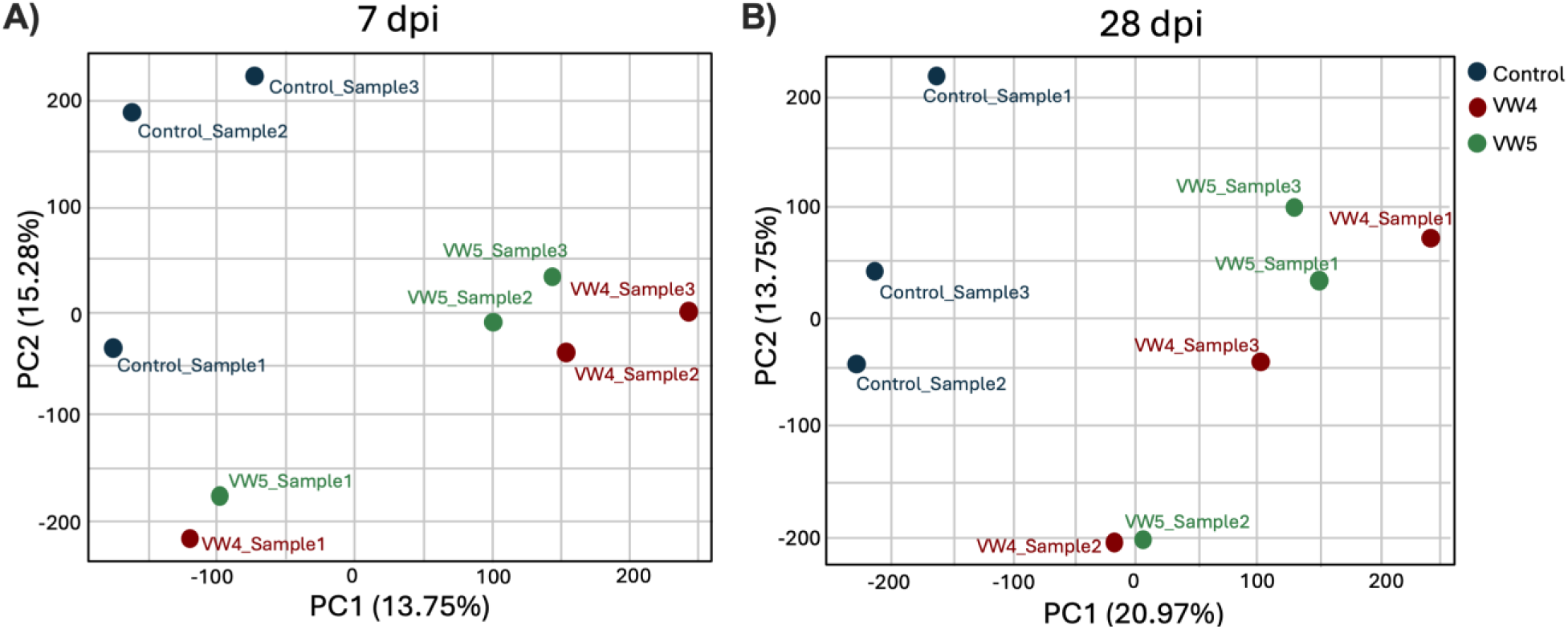
Principal component analysis distinguishes infected from uninfected cucumber roots at early and late infection stages. Principal component analysis (PCA) of mRNA-seq data shows clear separation between uninfected (control) cucumber roots and roots infected with *M. javanica* VW4 or VW5 at both 7 (A) and 28 dpi (B). The x and y axes show Principle Component 1 and Principle Component 2, which explains 20.97% and 15.2% of the total variance respectively. Each data point represents an independent biological replicate of RNA extracted from galled tissue of cucumber roots infected with VW4 or VW5, or from uninoculated control roots. Red data points indicate uninoculated control roots, green indicates VW4-infected roots, and blue indicates VW5-infected roots.

At 7 dpi, VW4 infection resulted in extensive transcriptional reprogramming, with 1,515 genes significantly upregulated and 981 genes downregulated relative to uninoculated controls (adjusted p-value < 0.05; log_2_(fold change) > 1; **Supp. Data 1-2 and Supp. Figure 6**). In contrast, VW5 infection at the same time point resulted in only 924 upregulated and 580 downregulated genes, indicating a markedly reduced change in host transcriptome response (adjusted p-value < 0.05; log_2_(fold change) > 1); **Supp. Data 3-4 and Suppl. Figure 6**). Removal of overlapping differentially expressed genes revealed that VW4 regulated a substantially larger gene set, with 709 uniquely upregulated and 564 uniquely downregulated genes. In comparison, VW5 uniquely regulated only 108 upregulated and 163 downregulated genes (**Figures 9A** and **9B**; (**Supp. Data 5-8**). Together, these results demonstrated that VW5 triggered a substantially weaker host transcriptional reprogramming during early infection compared with VW4.

**Figure 9:**
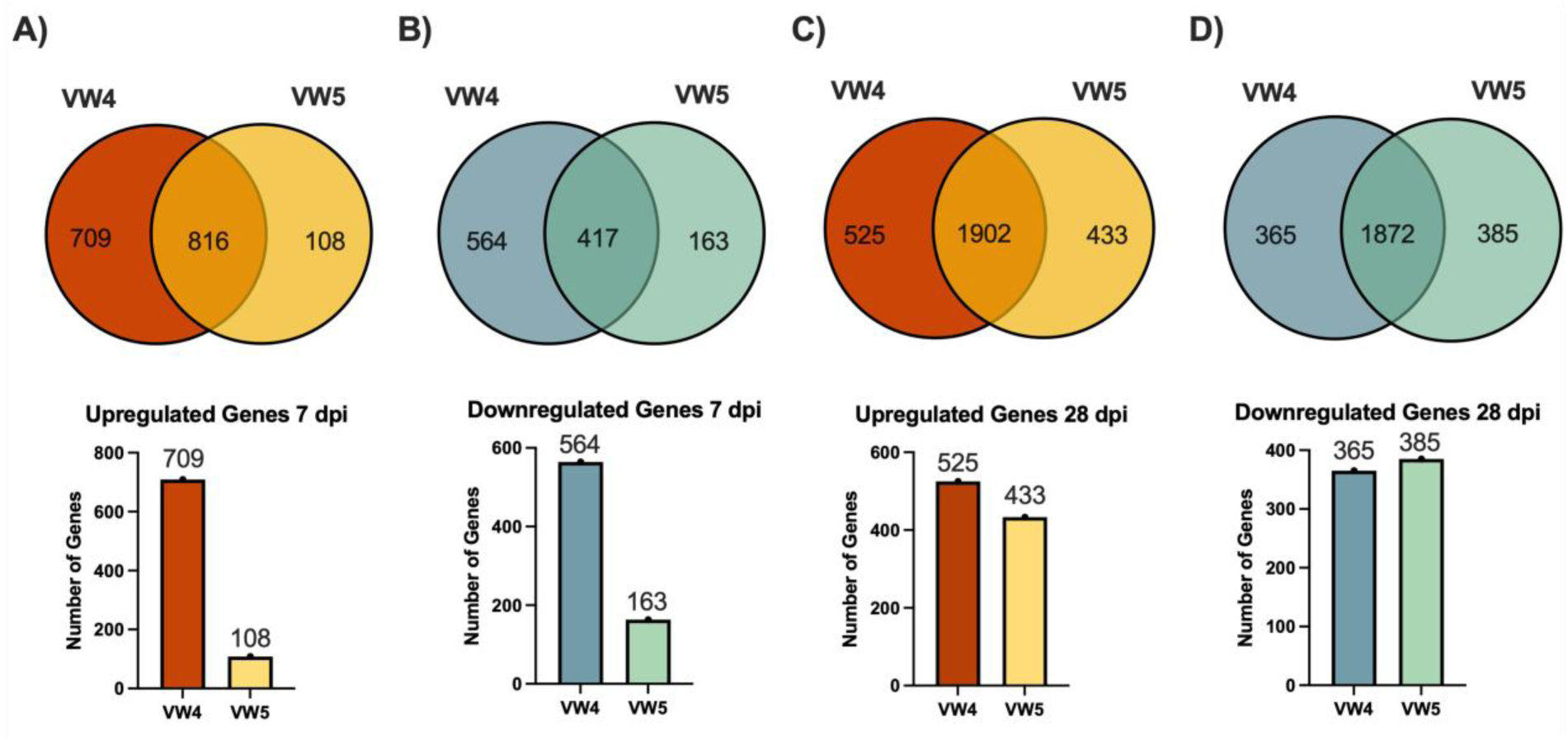
Differentially expressed cucumber genes unique to VW4 or VW5 show greater divergence during early infection. Venn diagrams showing significantly upregulated genes at 7 dpi (A), significantly downregulated genes at 7 dpi (B), significantly upregulated genes at 28 dpi (C), and significantly downregulated genes at 28 dpi (D). Numbers indicate genes uniquely regulated in either VW4 or VW5, while overlapping regions represent genes significantly regulated in both. Bar graphs summarizing the number of uniquely up- and downregulated genes in VW4 and VW5 at 7 and 28 dpi. Cool colors represent downregulated genes, and warm colors represent upregulated genes.

By 28 dpi, the overall magnitude of transcriptional change increased for both strains. VW4 infection resulted in 2,427 upregulated and 2,237 downregulated genes, while VW5 infection resulted in 2,335 upregulated and 2,257 downregulated genes (**Supp. Figure 6**). Analysis of uniquely regulated genes at this time point showed comparable numbers of strain-specific genes between VW4 and VW5, including 365 uniquely downregulated genes in VW4 compared to 385 in VW5, and 525 uniquely upregulated genes in VW4 compared to 433 in VW5 (**Figures 9C** and **9D; Supp. Data 9-12**). Notably, the number of overlapping differentially expressed genes was markedly higher at 28 dpi than at 7 dpi, indicating convergence of host transcriptional responses between the two strains during progress of parasitic relationship. Volcano plots displaying differentially expressed genes for VW4 and VW5 compared to control roots at both timepoints along with the top 5 differentially expressed genes are displayed in **Supp. Figure 7**.

To identify biological processes associated with strain-specific differences, we performed a gene ontology (GO) enrichment analysis using uniquely regulated genes by VW4 or VW5 at each time point (**Figures 10** and **11**). Because this analysis is restricted to unique genes, enriched GO terms reflect differences in nematode strain-specific regulation rather than the full host transcriptional response to nematode infection.

**Figure 10:**
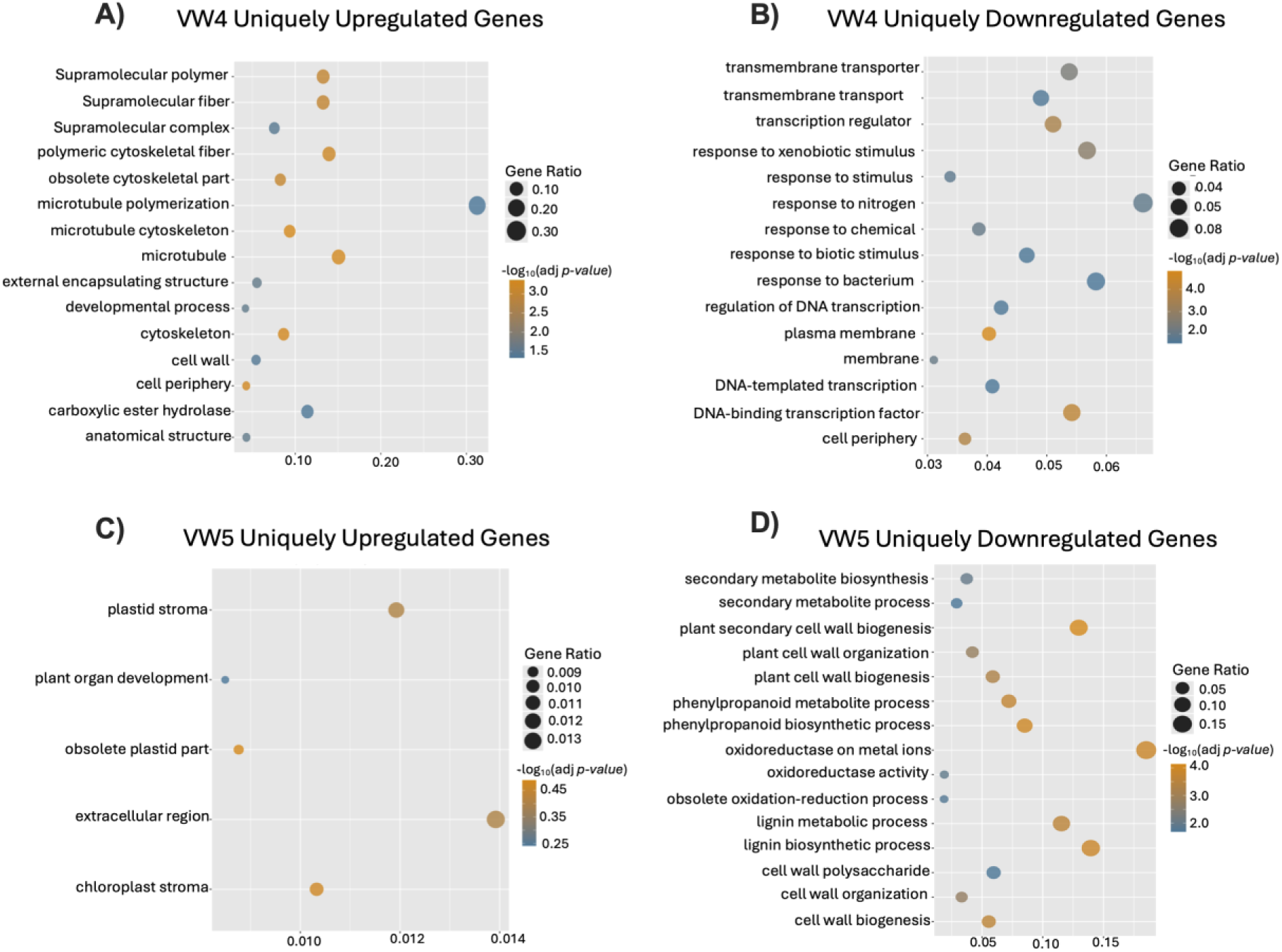
GO enrichment analysis of VW4 and VW5 uniquely differently expressed cucumber genes at 7 dpi. GO enrichment analysis was performed using genes uniquely up- or downregulated in cucumber roots infected with VW4 or VW5 strain of *M. javanica* at 7 dpi, relative to each other. Only genes that were significantly regulated respective to VW4 or VW5 were included in the analysis. VW4-uniquely upregulated genes (A), VW4-uniquely downregulated genes (B), VW5-uniquely upregulated genes (C), and VW5-uniquely downregulated genes (D). GO biological process terms are shown on the y-axis, and gene ratio is shown on the x-axis. Circle size represents the number of genes associated with each GO term (condensed to fit on chart), and circle color indicates significance based on −log10(adjusted p-value). The top 15 most significantly enriched GO terms are displayed.

**Figure 11:**
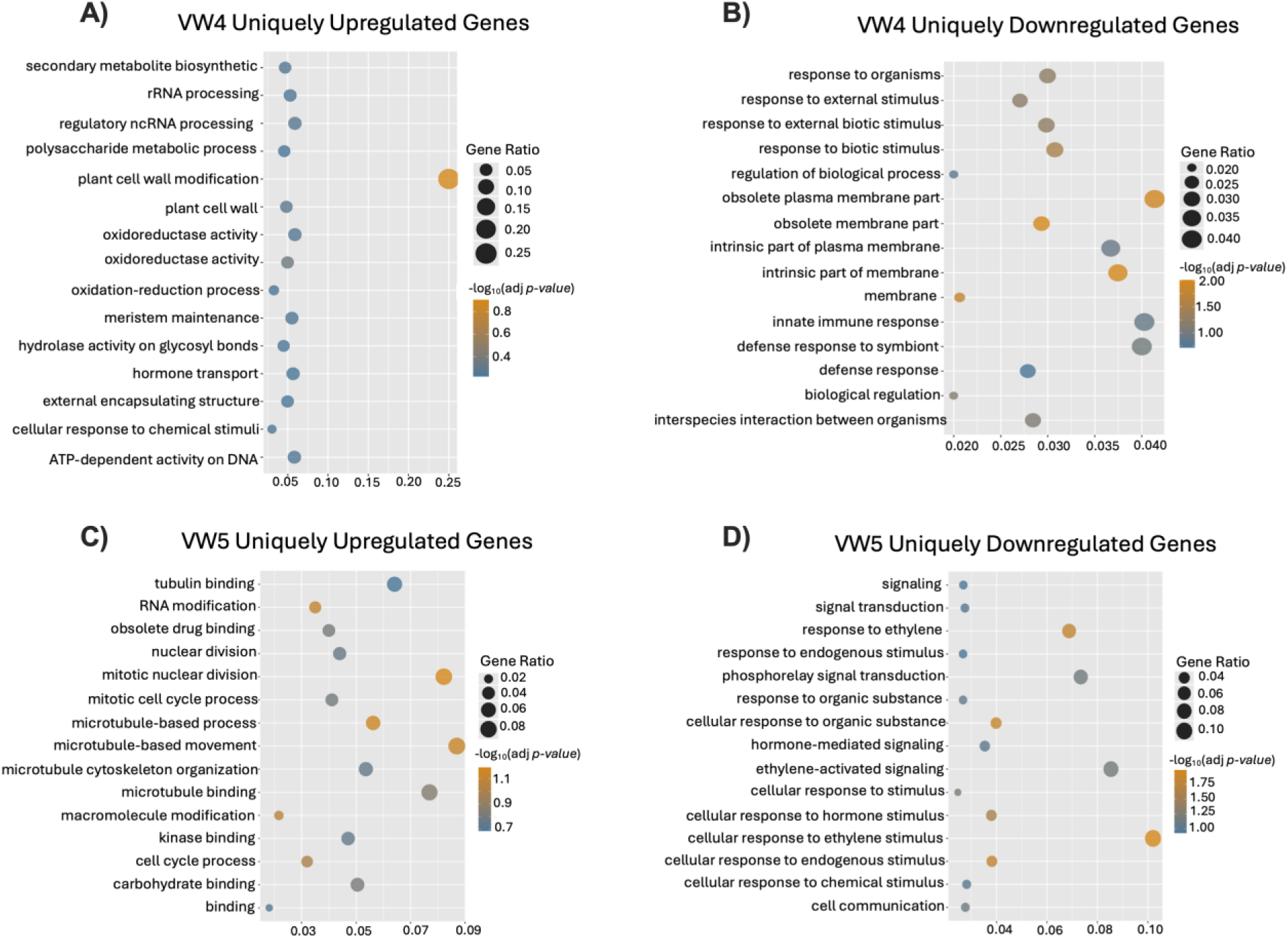
GO enrichment analysis of VW4-uniquely differentially expressed cucumber genes at 28 dpi. GO enrichment analysis was performed using genes uniquely up- or downregulated in cucumber roots infected with VW4 and VW5 strain of *M. javanica* at 28 dpi, relative to each other. Only genes that were significantly regulated respective to VW4 or VW5 were included in the analysis. VW4-uniquely upregulated genes (A), VW4-uniquely downregulated genes (B), VW5-uniquely upregulated genes (C), and VW5-uniquely downregulated genes (D). GO biological process terms are shown on the y-axis, and gene ratio is shown on the x-axis. Circle size represents the number of genes associated with each GO term (condensed to fit on chart), and circle color indicates significance based on −log10(adjusted p-value). The top 15 most significantly enriched GO terms are displayed.

At 7 dpi, VW4-uniquely regulated genes were significantly enriched for GO terms representing biological processes required for feeding site formation, including cellular reorganization, vascular development, cell cycle related processes, and modulation of host defense-related pathways (**Figures 10A and 10B; Supp. Figure 7A**) (Bartlem et al., 2014; de Almeida Engler et al., 2012). In contrast, VW5-unique genes at 7 dpi showed limited enrichment for feeding site–associated processes and were primarily enriched for secondary metabolic and plastid-related functions, indicating an impaired ability to reprogram host tissues morphogenesis during early infection (**Figures 10C** and **10D**; **Supp. Figure 7B**).

At 28 dpi, VW4-uniquely upregulated genes were enriched for processes associated with feeding site maintenance, cell wall regulation and metabolic activity, including RNA processing, hormone transport, meristem maintenance, and ATP-dependent activities (**Figure 11A** and **Supp. Figure 7C**). Whereas, VW4-uniquely downregulated genes were enriched for defense- and stimulus-response biological processes, indicating a downregulation of host immune-related pathways at this stage of infection (**Figure 11B** and **Supp. Figure 7D**). VW5-uniquely upregulated genes at 28 dpi were also enriched for processes associated with feeding site function and hormone-response pathways; however, many of these processes correspond to those enriched among VW4-unique genes at 7 dpi, suggesting a temporal delay in the activation of feeding site–associated pathways during VW5 infection (**Figures 10A** and **10B** and **Figures 11A** and **11C** and **Supp. Figure 7**). In contrast to VW4, VW5-uniquely downregulated genes did not show enrichment for defense-related GO terms, indicating limited suppression of defense-related processes at 28 dpi.

## Discussion

In this study, we demonstrate that Mi-1 resistance-breaking in *M. javanica* is associated with a measurable fitness cost on susceptible hosts. Specifically, the resistant-breaking strain VW5 produced fewer eggs than its avirulent progenitor VW4 on three susceptible hosts (tomato, cucumber, and rice). Importantly, this reduction in reproduction was not attributable to impaired root entry, as both strains entered host roots at comparable rates and reached similar total nematode numbers within roots. Instead, our results indicate that resistance breaking in VW5 compromises the nematode’s ability to establish and maintain functional feeding sites.

Microscopic analyses revealed abnormalities in gall morphology and GC structure in VW5-infected roots. VW4 induced the typical sequence of feeding site initiation, host cell reprogramming, and GC maturation characteristics of successful RKN parasitism, whereas VW5-induced GCs were structurally compromised during early infection. These defects included reduced cellular hypertrophy, altered GC organization, limited vascular connectivity, and poorly developed cell wall ingrowths. The phenotype was particularly pronounced in cucumber, consistent with the stronger reduction in egg production observed on this host. Because giant cells are essential for sustained nutrient acquisition, these structural defects likely underlie the reduced fecundity of VW5. Transcriptomic analyses further support this interpretation, indicating that VW5 induced a narrower set of host transcriptional changes than VW4, particularly within gene categories associated with feeding site initiation and host cell reprogramming. Importantly, because GO enrichment was restricted to strain-specific gene sets, these differences do not indicate a complete absence of host manipulation by VW5 but rather reflects the loss of specific components of the transcriptional reprograming program deployed by VW4.

Based on these findings, we propose a trade-off model in which acquisition of Mi-1 virulence involves loss or alteration of a nematode avirulence (Avr) gene product that is required both for recognition by the Mi-1 resistance pathway and for optimal host manipulation. Loss of such an Avr gene would enable evasion of Mi-1–mediated defense while simultaneously compromising feeding site establishment, nutrient acquisition, and reproduction on susceptible hosts. This hypothesis is consistent with previous work demonstrating that VW5 arose directly from VW4 under selection on Mi-1 tomato and that the two strains are nearly genetically identical, differing primarily by the absence of the Cg-1 cDNA fragment required for Mi-1–mediated resistance (Gleason et al., 2008). However, given the polyploid and asexual nature of RKN genomes, resistance breaking in VW5 may involve additional gene losses, mutations, or structural. Resolving the full genetic basis of this trade-off will require haplotype-resolved genome comparisons between VW4 and VW5.

Although our results are consistent with a loss-of-Avr trade-off model, alternative evolutionary scenarios are also possible. Resistance breaking could, in some cases, arise through gain-of-function changes that enhance suppression of host immune signaling pathways rather than through loss of an Avr gene. For example, mutations that increase the efficacy or expression of effectors targeting defense-associated signaling networks could permit evasion of Mi-1–mediated recognition while maintaining or even enhancing host manipulation capacity. Under such a model, resistance breaking would not necessarily incur a fitness penalty and could explain the robust reproductive performance observed in some independently evolved field populations (Iberkleid et al., 2014; Ploeg et al., 2023;). Distinguishing between loss-of-function and gain-of-function mechanisms will require functional characterization of candidate effectors and comparative genomic analyses of multiple virulent lineages.

From an applied perspective, these results underscore both the vulnerability and resilience of single-gene resistance strategies. Although Mi-1 has remained effective for decades, the emergence of resistance-breaking populations combined with environmental pressures such as increasing soil temperatures raises concerns regarding durability. However, the reduced performance of virulent strains on susceptible hosts suggests that resistance breaking may not be selectively advantageous in the absence of Mi-1–mediated selection. Such trade-offs could slow the spread or dominance of virulent genotypes in mixed cropping systems or rotations incorporating susceptible cultivars. Determining how frequently such fitness costs occur in field populations will be critical for predicting long-term resistance durability.

Finally, the VW4-VW5 system provides a powerful framework for identifying nematode effectors and host susceptibility pathways that underpin successful parasitism. Genes and pathways identified through comparative genomic and transcriptomic analyses genomic comparisons may serve as targets for next-generation control strategies, including susceptibility gene editing or diagnostic markers for early nematode infection. Together, our results demonstrate that resistance breaking in root-knot nematodes is not cost-free and that dissecting these trade-offs offers valuable insight into both the evolution of virulence and the development of durable nematode management strategies.

