## Supplementary Table 1 for "Resistance Breaking in Root-knot Nematodes Carries a Fitness Cost Associated with Defective Feeding Site Development"

**Supplemental Figures**

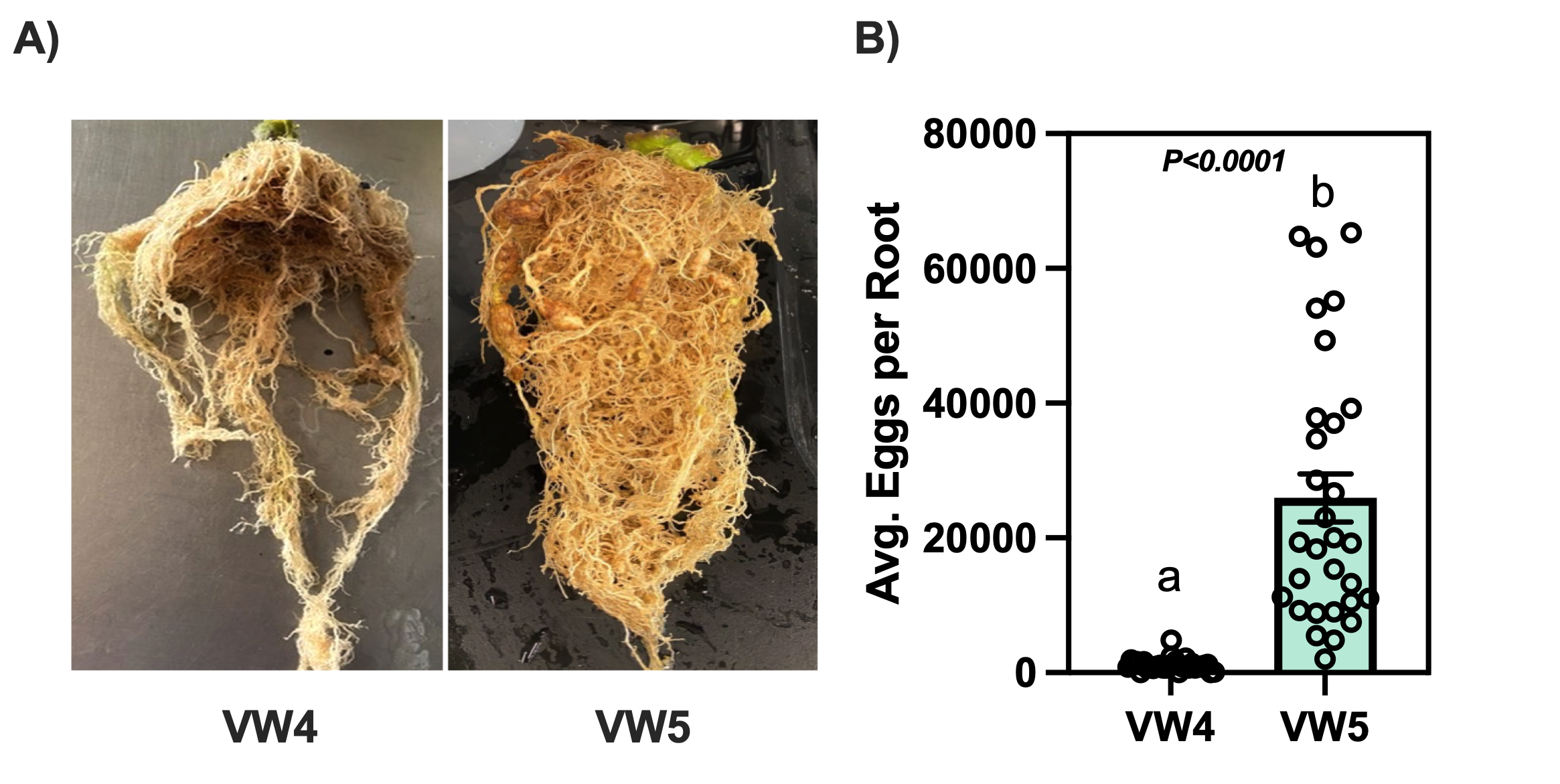

**Supplementary Figure 1: A near isogenic population of *M. javanica* (VW5) overcomes resistance of *Mi-1* on tomato.** (A) Resistant tomato (cv. Motelle) containing the *Mi-1* gene infected with *M. javanica* wildtype VW4 and resistance-breaking VW5. (B) The average numbers of eggs produced by VW5 and VW4 on cv. Motelle. Each data point represents an individual root system. Bars indicate means ± standard error of the mean (SEM) Statistical significance was assessed using an unpaired Student’s t-test; significant difference is indicated by letters, (n = 30 roots per treatment).

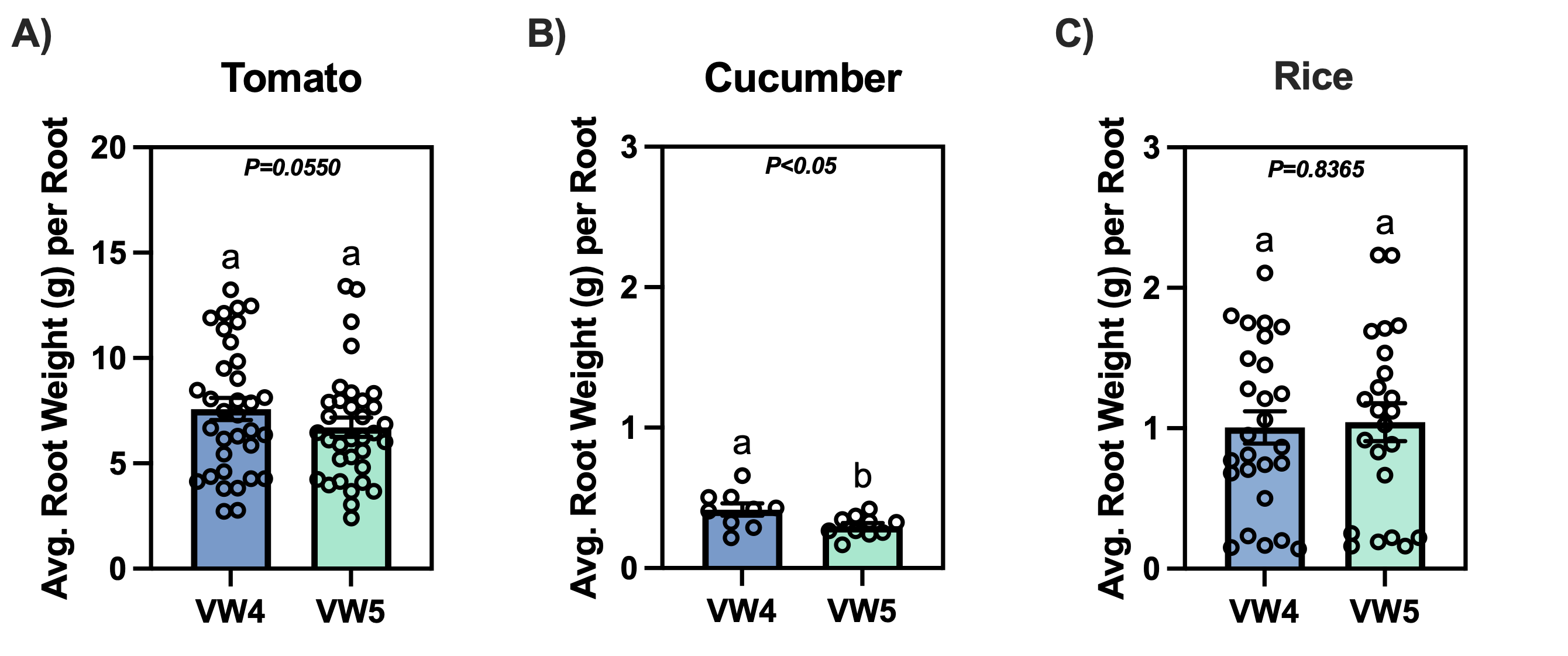

**Supplementary Figure 2: Root fresh weights show a difference between roots infected with VW4 or VW5 for cucumber only**. (A) Average tomato root weight, (B) Average cucumber root weight, (C) Average rice root weight. Fresh weight is recoded in grams. Each data point represents an individual root system. Bars indicate means ± standard error of the mean (SEM). Statistical significance was determined using unpaired Student’s t-tests, with p values shown; different letters indicate statistically significant differences (p < 0.05). Data are pooled from three independent biological experiments (tomato, n = 34; cucumber, n = 18; rice, n = 26).

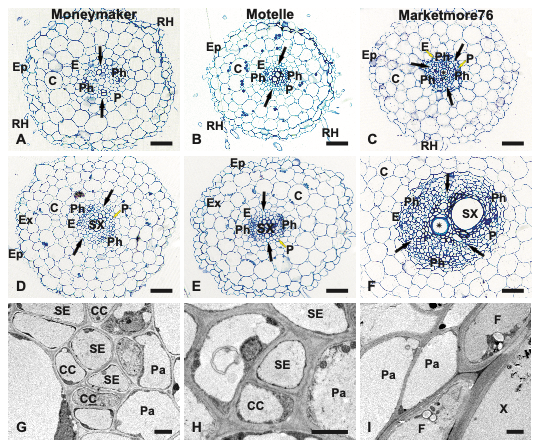

**Supplementary Figure 3: Anatomy of uninfected roots of tomato and cucumber.** Images of traversed sections of uninfected tomato roots of susceptible cv. Moneymaker (A and D), resistant cv. Motelle (B and E) and susceptible cucumber cv. Marketmore76 (C and F) at the late stage of primary (A-C) and the early stage of secondary root growth (D-F). Abbreviations: C, cortex; Ep, epidermis; Ex, exodermis; P, pericycle; Ph, phloem; RH, root hair; SX, secondary xylem vessel. Small yellow arrows indicate positions of tissues. Black arrows indicate the axis of primary xylem bundles. Asterisk indicates central metaxylem vessel. Scale bars: 50 µm.

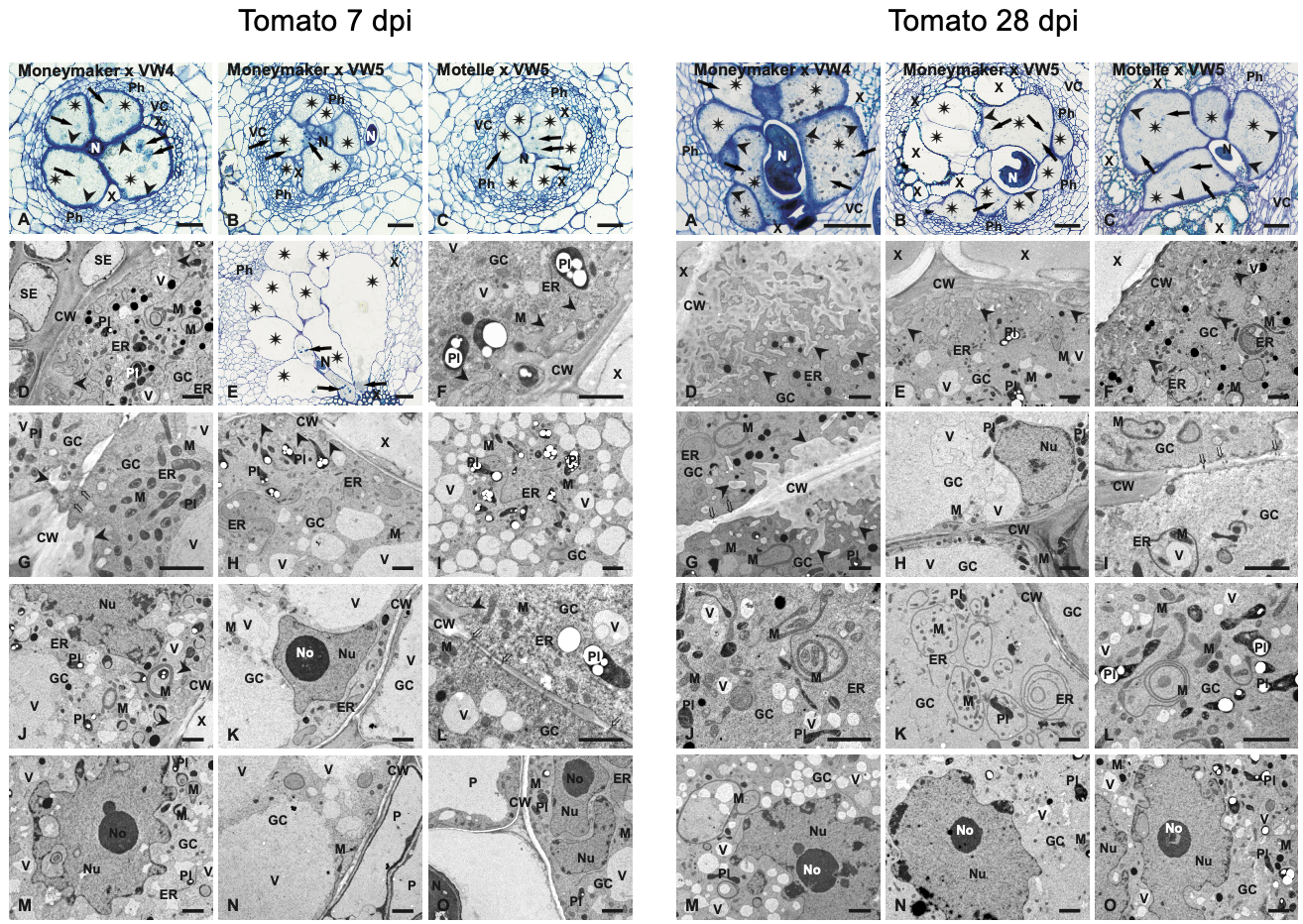

**Supplemental Figure 4: Anatomy and ultrastructure of giant cells developed in tomato roots at 7 dpi.** Light (A-C and E) and transmission electron microscopy (D and F-O) images of transverse sections of susceptible cv. Moneymaker (A, B, D, E, G, H, J, K, M and N) and resistant cv. Motelle (C, F, I, L, and O) infected with avirulent VW4 strain (A, D, G, J and M) or virulent VW5 strain (B, C, E, F, H, I, K, L, N and O). Asterisks indicate giant cells (A-C and E). Arrows point to giant cell nuclei (A-C and E). Arrowheads indicate systems of cell wall ingrowths (A, D, F, G, H, J and L). Double tails arrows indicate plasmodesmata in thin fragments of cell walls between giant cells (G, I, and L). Scale bars: 50 µm (A-C and E) and 2 µm (D and F-O). Abbreviations: CW, cell wall; ER, endoplasmic reticulum; GC, giant cell; M, mitochondrion; N, nematode; No, nucleolus; Nu, nucleus; P, parenchymatic cell; Ph, phloem; Pl, plastid; SE, sieve tube element; X, xylem vessel; V, vacuole.

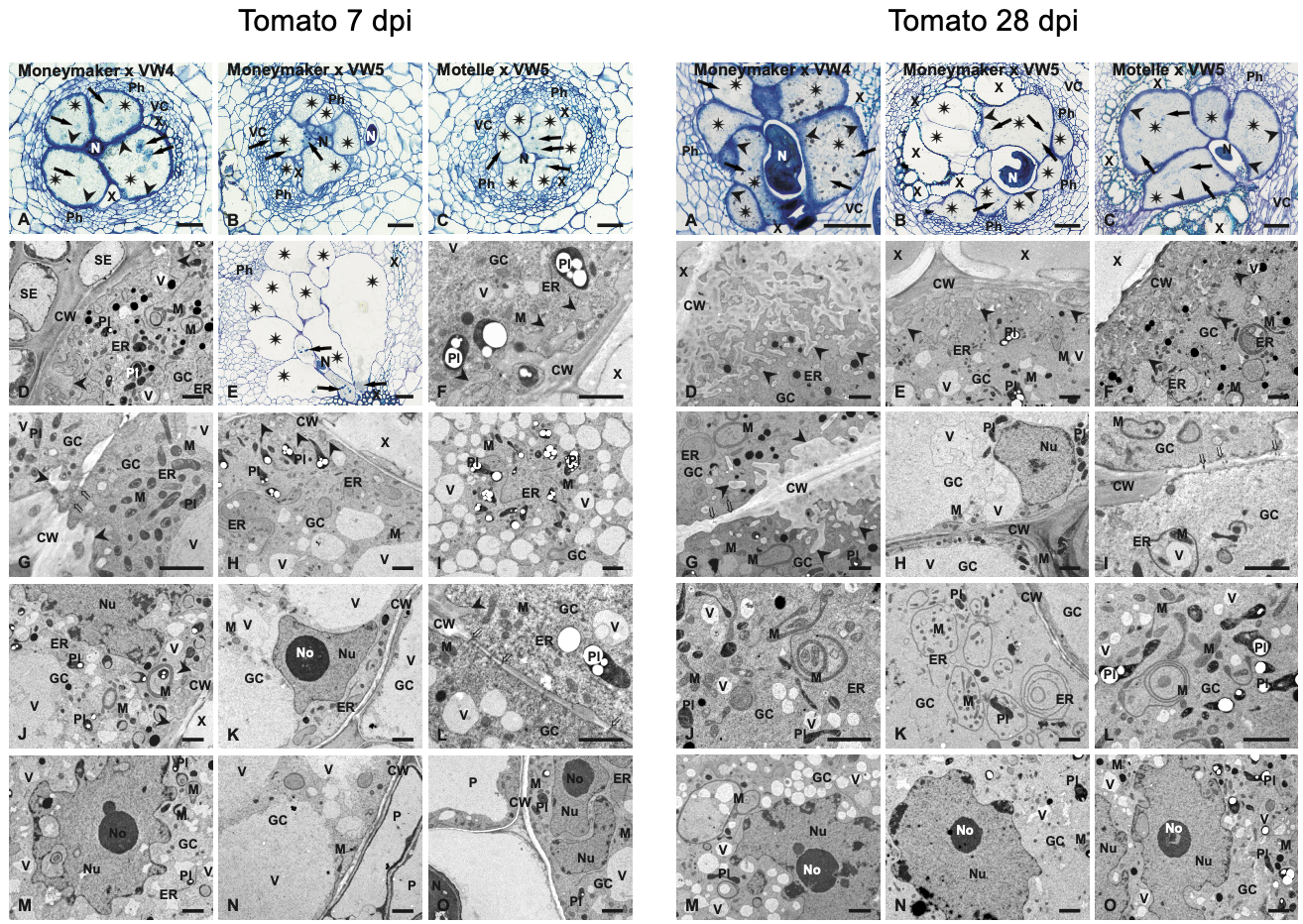

**Supplementary Figure 5: Anatomy and ultrastructure of giant cells developed in tomato roots at 28 dpi.** Light (A-C) and transmission electron microscopy (D-O) images of transverse sections of susceptible cv. Moneymaker (A, B, D, E, G, H, J, K, M and N) and resistant cv. Motelle (C, F, I, L, and O) infected with avirulent VW4 strain (A, D, G, J and M) or virulent VW5 strain (B, C, E, F, H, I, K, L, N and O). Scale bars: 50 µm (A-C) and 2 µm (D-O). Abbreviations: CW, cell wall; ER, endoplasmic reticulum; GC, giant cell; M, mitochondrion; N, nematode; No, nucleolus; Nu, nucleus; P, parenchymatic cell; Ph, phloem; Pl, plastid; X, xylem vessel; V, vacuole.

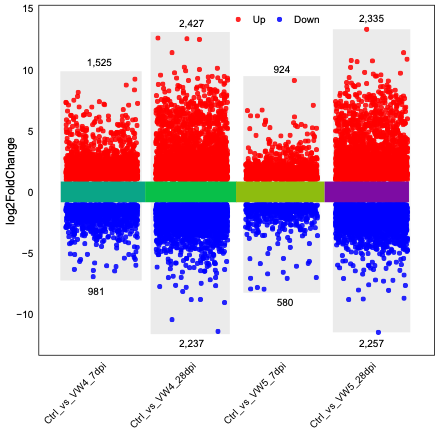

**Supplementary Figure 6: Rplot representation of all significantly up- and downregulated genes compared to uninoculated roots at 7 and 28 dpi.** Cucumber roots were inoculated with VW4 or VW5 strain of *M. javanica.* Galls were collected at 7 and 28 dpi. Gene significance was compared individually to the respective control roots. Significantly upregulated genes are shown in red and significantly downregulated genes are shown in blue. The number of genes is listed under and above each set. y-axis shows log2_foldchange.

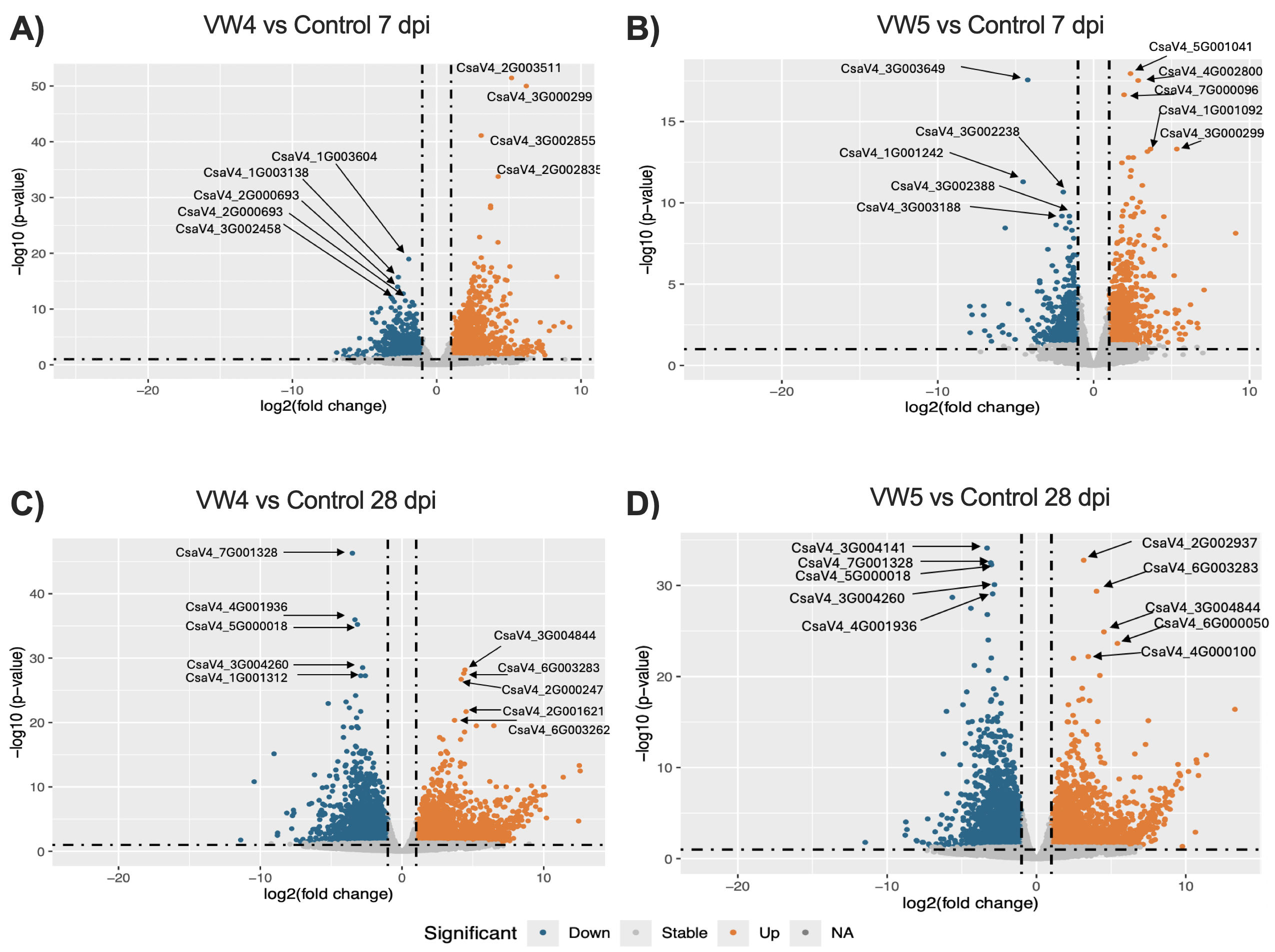

**Supplementary Figure 7: Cucumber genes are more significantly and abundantly regulated during VW4 than VW5 infection.** Volcano plots were generated using ggplot2 package in R Studio and shows genes expressed in cucumber during infection with either VW4 or VW5 at 7 and 28 dpi. (A) Significant gene expression after VW4 infection compared to the uninoculated roots at 7dpi. (B) Significant gene expression after VW5 infection compared to the uninoculated roots at 7 dpi. (C) Significant gene expression after VW4 infection compared to uninoculated roots at 28 dpi. (D) Significant gene expression after VW5 infection compared to uninoculated roots at 28 dpi. p < 0.05 for significance cut off. Significantly upregulated genes are shown in orange, while significantly downregulated genes are shown in blue, non-significant genes are grey (behind and under the dotted lines). The x-axis represents the log_2_(foldchange) and the y-axis represents the -log10(adjusted p-value).

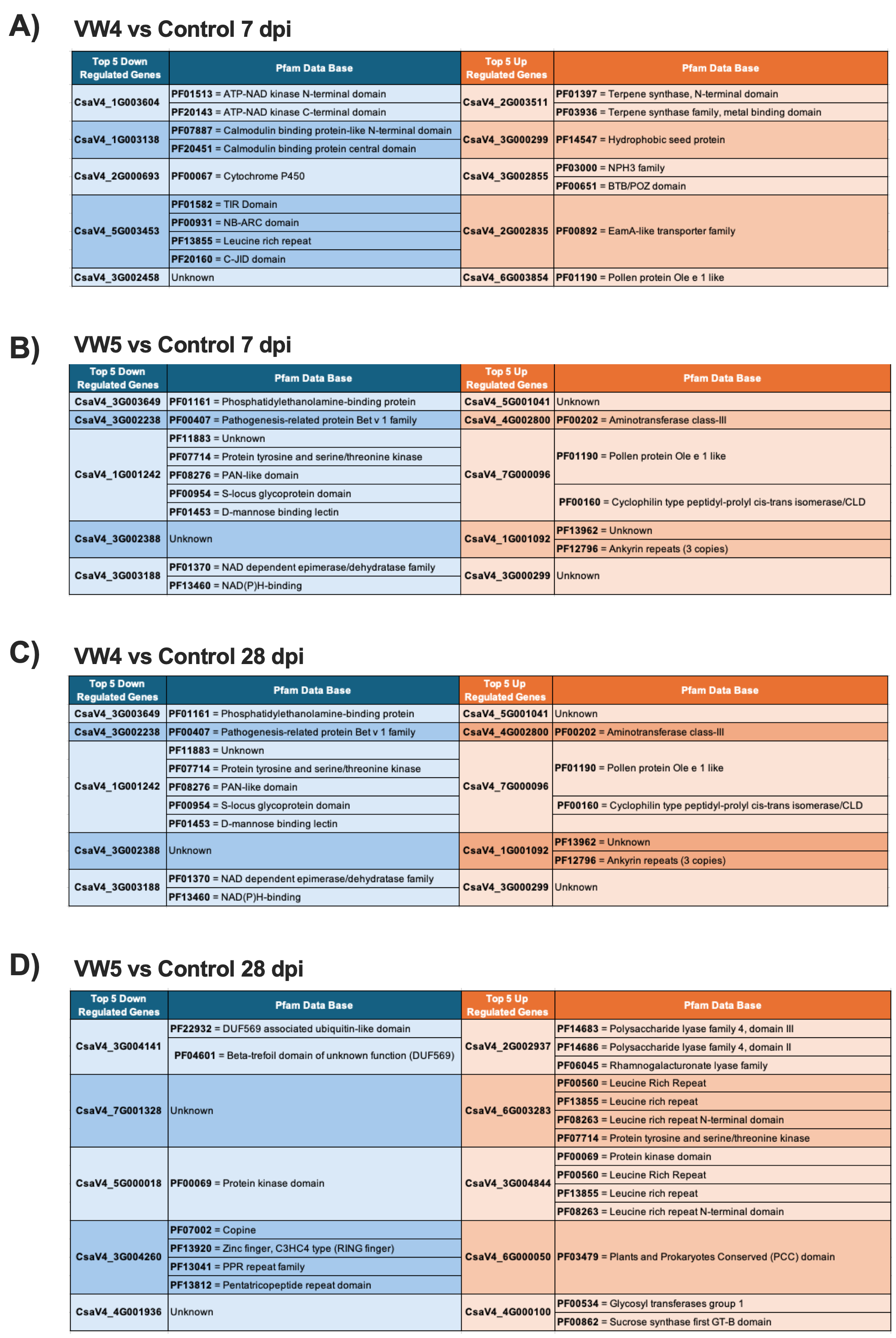

**Supplementary Figure 8: Top 5 most significantly expressed up- and downregulated genes functions and domains.** Top 5 most significant downregulated genes in blue. Top 5 most significant upregulated in orange. (A) Top 5 significant genes up- and downregulated for VW4 infection at 7 dpi, (B) Top 5 significant genes up- and downregulated for VW5 infection at 7 dpi, (C) Top 5 most significant genes up- and downregulated for VW4 infection at 28 dpi, and (D) Top 5 most significant genes up and downregulated for VW5 infection at 28 dpi. Gene number is listed along with associated Pfam predicted domain and function for each gene using InterPro data base.

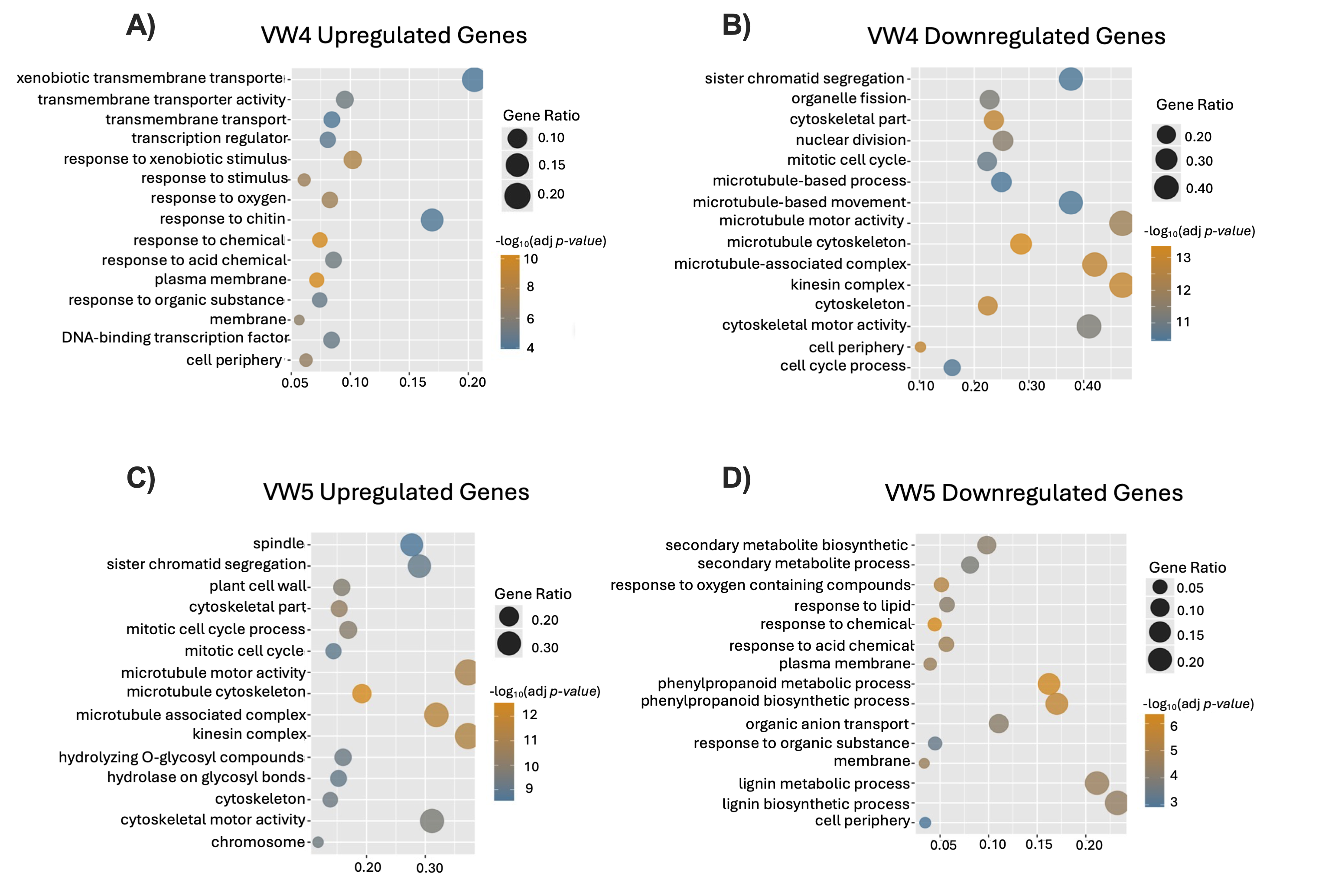

**Supplementary Figure 9**: **GO-enrichment analysis of total cucumber genes up- and downregulated genes at 7 dpi.** (A) Significantly upregulated gene families for VW4 infected roots at 7 dpi. (B) Significantly downregulated gene families for VW4 infected roots at 7 dpi. (C) Significantly upregulated gene families following VW5 infected roots at 7 dpi. (D) Significantly downregulated gene families following VW5 infected roots at 7 dpi. GO biological process terms are shown on the y-axis, and gene ratio is shown on the x-axis. Size of the circle indicates number of genes associated with each function. Circle color indicates p-value significance where -log10(adjusted p-value) and p<0.05. Top 15 genes are displayed.

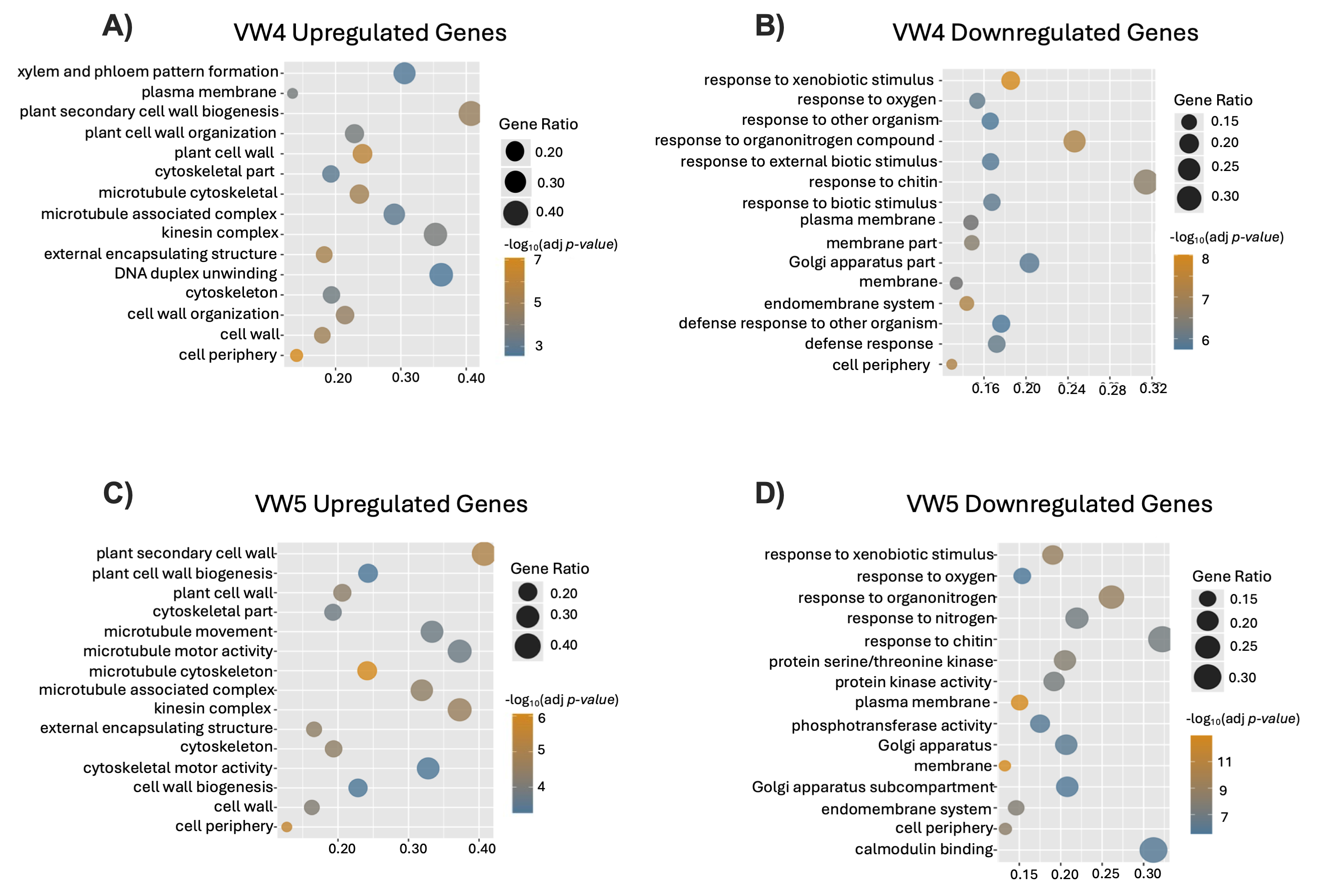

**Supplementary Figure 10: GO-enrichment analysis of total cucumber genes up- and downregulated genes at 28 dpi.** (A) Significantly upregulated gene families for VW4 infected roots at 28 dpi. (B) Significantly downregulated gene families for VW4 infected roots at 28 dpi. (C) Significantly upregulated gene families following VW5 infected roots at 28 dpi. (D) Significantly downregulated gene families following VW5 infected roots at 28 dpi. GO biological process terms are shown on the y-axis, and gene ratio is shown on the x-axis. Circle size indicates number of genes associated with each function. Circle color indicated p-value significance where -log10(adjusted p-value) and p<0.05. Top 15 genes are displayed.
